# A domestication history of dynamic adaptation and genomic deterioration in sorghum

**DOI:** 10.1101/336503

**Authors:** Oliver Smith, William V Nicholson, Logan Kistler, Emma Mace, Alan Clapham, Pamela Rose, Chris Stevens, Roselyn Ware, Siva Samavedam, Guy Barker, David Jordan, Dorian Q Fuller, Robin G Allaby

**Affiliations:** School of Life Sciences, University of Warwick, Coventry, CV4 7AL, United Kingdom; Natural History Museum of Denmark, Øster Voldgade 5-7, 1350 København K, Denmark; Warwick Medical School, University of Warwick, Coventry, CV4 7AL, United Kingdom; Department of Anthropology, Smithsonian Institution, National Museum of Natural History, Washington, D.C. 20506, USA; Department of Agriculture, Fisheries and Forestry Queensland (DAFFQ), Warwick, Queensland 4370, Australia; The Austrian Archaeological Institute, Cairo Branch, Zamalek, Cairo, Egypt; Institute of Archaeology, UCL, London, United Kingdom; Queensland Alliance for Agriculture and Food Innovation, The University of Queensland, Warwick, Queensland 4370, Australia

**Keywords:** Ancient DNA, archaeobotany, bottleneck, introgression, genomic rescue

## Abstract

The evolution of domesticated cereals was a complex interaction of shifting selection pressures and repeated introgressions. Genomes of archaeological crops have the potential to reveal these dynamics without being obscured by recent breeding or introgression. We report a temporal series of archaeogenomes of the crop sorghum (*Sorghum bicolor*) from a single locality in Egyptian Nubia. These data indicate no evidence for the effects of a domestication bottleneck but instead suggest a steady decline in genetic diversity over time coupled with an accumulating mutation load. Dynamic selection pressures acted sequentially on architectural and nutritional domestication traits, and adaptation to the local environment. Later introgression between sorghum races allowed exchange of adaptive traits and achieved mutual genomic rescue through an ameliorated mutation load. These results reveal a model of domestication in which genomic adaptation and deterioration was not focused on the initial stages of domestication but occurred throughout the history of cultivation.

---

The evolution of domesticated plant forms represents a major transition in human history that facilitated the rise of modern civilization. In recent years our understanding of the domestication process has become revised considerably (1). In the case of cereals it has been recognized that the selective forces that give rise to domestication syndrome traits such as the loss of seed shattering were generally weak and comparable to natural selection (2,3) and that the intensity of selection pressures changed over the course of time as human technology evolved (4). Furthermore, domesticated lineages have often been subjected to repeated introgressions from local wild populations that endowed adaptive traits and obscured historical signals in the genome (5). Such complexity obfuscates attempts to reconstruct the evolutionary history of domesticated species from modern plants. To counter these confounding factors in this study we directly tracked the evolutionary trajectory of a domesticated species, sorghum (*Sorghum bicolor* ssp. *bicolor* (L.) Moench.), through the archaeological record. This approach enabled the identification of selection pressures not clear today, and the tracking of the introgression process, revealing a domestication history which runs counter to the expectations of the current conventional model of domestication.

Sorghum is the world’s fifth most important cereal crop and the most important crop of arid zones (6) used for food, animal feed, fibre and fuel. The evolution of sorghum has seen its transition from being a wild pluvial plant in north-eastern Africa (*S. bicolor* ssp. *verticilliflorum* (Steud.) De Wet ex Wiersema & Dahlberg, hereafter referred to as *S. verticilliflorum* for clarity) to the ancestral domesticated form *Sorghum bicolor* type bicolor in Central Eastern Sudan by around 5000 years ago, while cultivation is inferred to have begun by 6000 yrs BP (7)., Ultimately, four specialized agroclimatic adapted types evolved after domestication—durra, caudatum, guinea, and kafir (8–10). The derived types were likely founded on introgressions of the wild progenitor complex *Sorghum verticilliflorum* or closely related species into the ancestral bicolor type, endowing traits such as drought tolerance in the case of type durra (8,11). The evolutionary history of sorghum, replete with introgression, is difficult to reconstruct from modern datasets. However, a temporal series of archaeobotanical domesticated sorghums spanning back to 2100 before present (yrs BP) at the archaeological site of Qasr Ibrim, situated on the Nubian frontier of northern Africa, affords the opportunity to track this complex crop directly through time removing the obscuring effects of introgression (12). Prior to this wild sorghum is present at Qasr Ibrim from at least ca. 2800 yrs BP. Domesticated sorghum (race bicolor) appears at the site ca. 2100 yrs BP. After this time period phenotypically domesticated sorghum of the ancestral type bicolor occurs throughout all cultural periods until the site was abandoned 200 years ago. During the early Christian period at 1470 yrs BP, the oldest known drought-adapted, free-threshing durra type appears at the site and occurs there for the rest of the site’s occupancy. The origins of the durra type are unclear. Current distributions in northern and eastern Africa and its dominance in the Near East and South Asia led to the proposal that durra originated on the Indian subcontinent (13) and returned to Africa at some point after 2000 yrs BP (14).

## Results

### Genetic diversity of sorghum over time

To gain a longitudinal insight into the evolutionary history of sorghum, we sequenced 9 archaeological genomes from different time points at Qasr Ibrim, including a wild phenotype from 1765 yrs BP and 8 domesticated phenotypes between 1805 and 450 yrs BP, a further 2 genomes from herbarium material, and 12 genomes of modern wild and cultivated sorghum types representing the varietal range (Table S1, S2).

We investigated how genetic diversity has changed through time by measuring within genome heterozygosity of 100 kbp genomic blocks that revealed a pattern of broad variation in heterozygosity in the wild progenitor *S. verticilliflorum* that became progressively narrower over time in the ancestral bicolor type, Figure S1. Interestingly, the wild phenotype of sorghum at Qasr Ibrim (sample A3) has a narrower variation in heterozygosity than the wild progenitor (represented by modern wild diversity), suggesting that it had been already been subject to genetic erosion. Conversely, the durra types all showed similar low levels of genomic variation in heterozygosity suggestive of genetic erosion prior to their appearance at Qasr Ibrim. Total genomic heterozygosity of bicolor over time confirmed that the ‘wild’ sorghum had already undergone considerable genetic erosion relative to the wild progenitor. To our surprise, the decreasing trend in heterozygosity over time fits a linear model (*p* values 0.0041 and 9.2×10^−6^ for parameters a and b respectively) better than an exponential model (*p* values 0.042 and 9.3×10^−5^) as would be expected from an early initial loss of diversity through a domestication bottleneck (15), Figure 1, Table S3 suggesting that there was no measurable effect on genetic diversity attributable to a domestication bottleneck.

**Figure 1.**
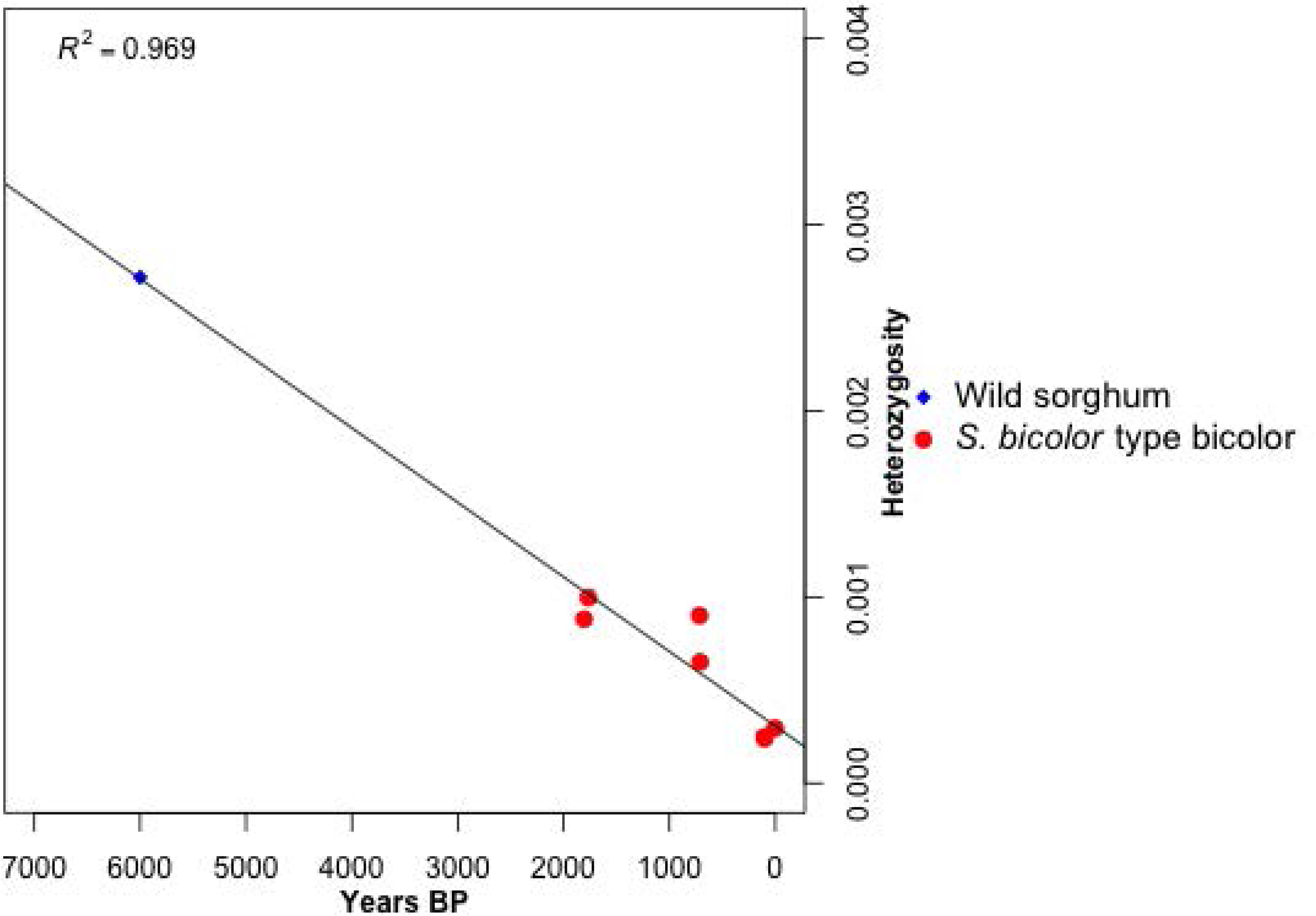
Genomic heterozygosity over time in *S. bicolor* type bicolor.

### Mutation load over time in Sorghum

The apparent lack of a domestication bottleneck runs contrary to expectations for a domesticated crop. To investigate the apparent lack of a domestication bottleneck further, we considered the mutation load. An expected consequence of the bottleneck is a rise in mutation load as small populations incorporate deleterious mutations through strong-acting drift. High mutation loads have generally been observed in domesticated crops (16–18), which have been taken as a confirmation of the effects of the domestication bottleneck. We measured the mutation load over time in the archaeological sorghum using a genome evolutionary rate profiling (GERP) analysis considering the total number of potentially deleterious alleles (19) (see methods), Figure 2. As with other domesticated crops, modern sorghum has a higher mutation load than its wild progenitor under both recessive and additive models. In contrast to the expectations of a domestication bottleneck we did not observe an initial large increase in mutation load associated with domestication, but rather an overall increasing trend in mutation load over time to the present day suggesting a process of load accumulation combined with selective purging episodes. In this case the trend line is best described by a positive exponential model rather than a linear model, Table S3, indicating mutation load has become increasingly problematic in recent times. However, the p-values suggest that the coefficient for the time in each model is only weakly significant, which could be the result of multiple processes on going, such as a strong increase in the rate of mutation load accumulation in recent times. When we considered the number of sites containing deleterious alleles (dominant model) rather than total number of deleterious alleles we observe a decreasing trend over time (Figure S3). This pattern suggests that part of the rising mutation load in the bicolor type was due to the increased homozygosity over time causing fixation of deleterious alleles originating from the wild progenitor pool. There is variation over time in mutation load, most notably in 1805 year-old sorghum (sample A5) that shows a sharp increase due to the incorporation of strongly deleterious alleles, both in the total number of alleles and number of sites. Interestingly, we found that the durra types show a pattern that contrasts to the bicolor type with relatively little change in heterozygosity and a significant fall in mutation load over time, suggesting the purging of deleterious mutations either through selection or genomic rescue through hybridization (Figure S2, S3). The contrasting patterns in mutation load over time are also reflected in methylation state profiles, which can reflect the state of genome-wide stress (20), Figure S4.

**Figure 2.**
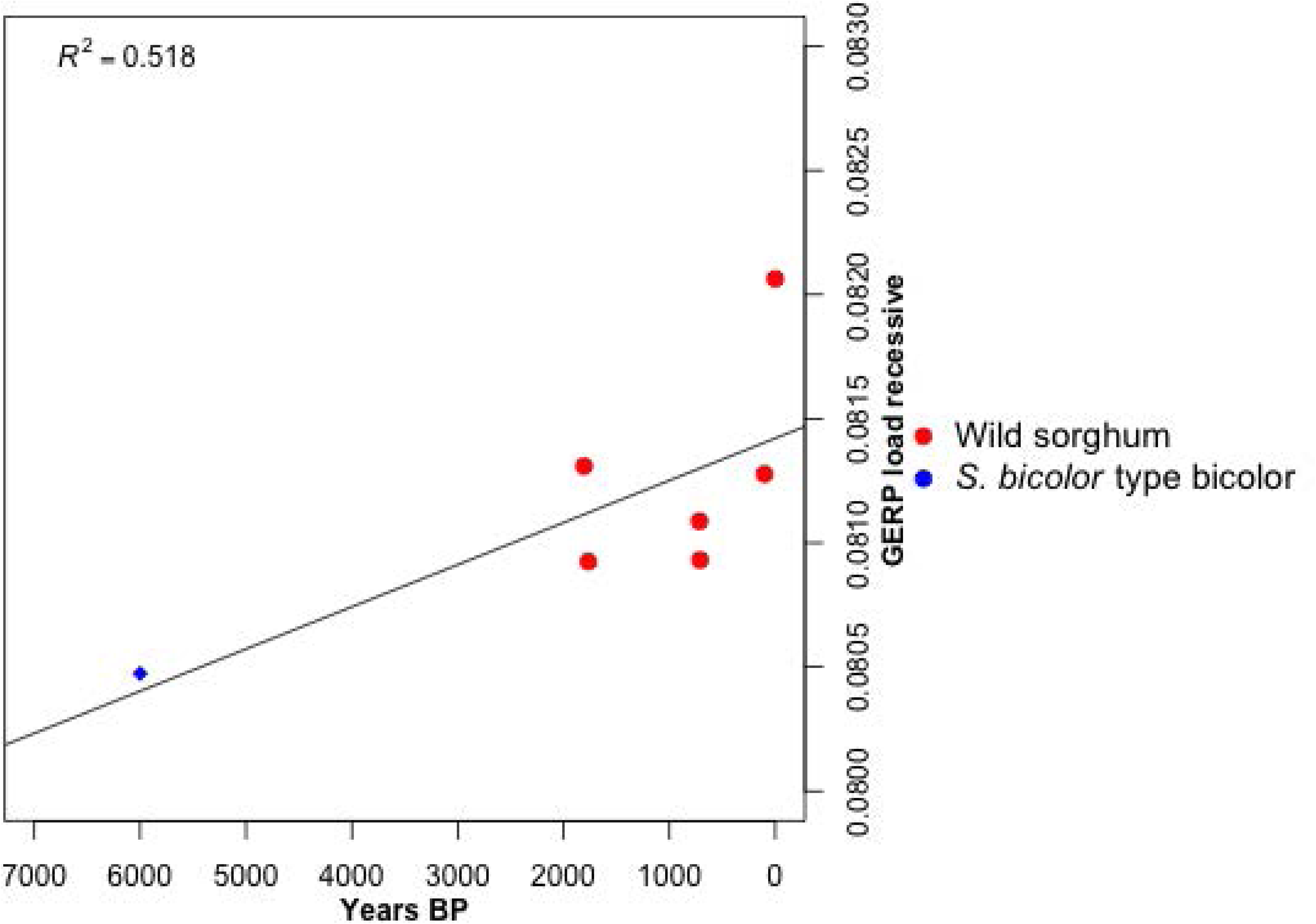
Total recessive GERP load over time in *S. bicolor* type bicolor.

### Signals of selection in sorghum

We considered that episodes of selection could have contributed in part to the variation in mutation load observed over time, either through reducing population size due to the substitution load or through hitchhiking effects. Three approaches were used to identify candidate regions under selection. Firstly, we surveyed for wild/domestication heterozygosity to look for significant reduction in heterozygosity in domesticates, which revealed 30 peaks of genome-wide significance (denoted by prefix pk), Figure 3, Table S4, S5. We also specifically surveyed 38 known domestication loci and also found a significant reduction of heterozygosity in 15 of the 38 associated regions, Table S6. Secondly, we used a SweeD analysis to detect selective sweeps (21), which identified 11 peaks (denoted by prefix s), Figure 3, Table S7. In the third approach we utilized the temporal sequence of archaeogenomes to investigate episodes of selection intensification by considering the gradient of heterozygosity change over time (see methods). In this latter approach we tracked the gradient of change in the heterozygosity of regions identified in the first two approaches and assigned significance based on the gradient deviation from the genome average for each type. We considered multiple time sequences representing alternative possible routes through contemporaneous genomes over time within type bicolor and type durra respectively. This revealed a period of selection intensification associated with domestication loci prior to 1805 yrs BP, followed by oscillations in diversity, Table S8, Figures S5–S7.

**Figure 3.**
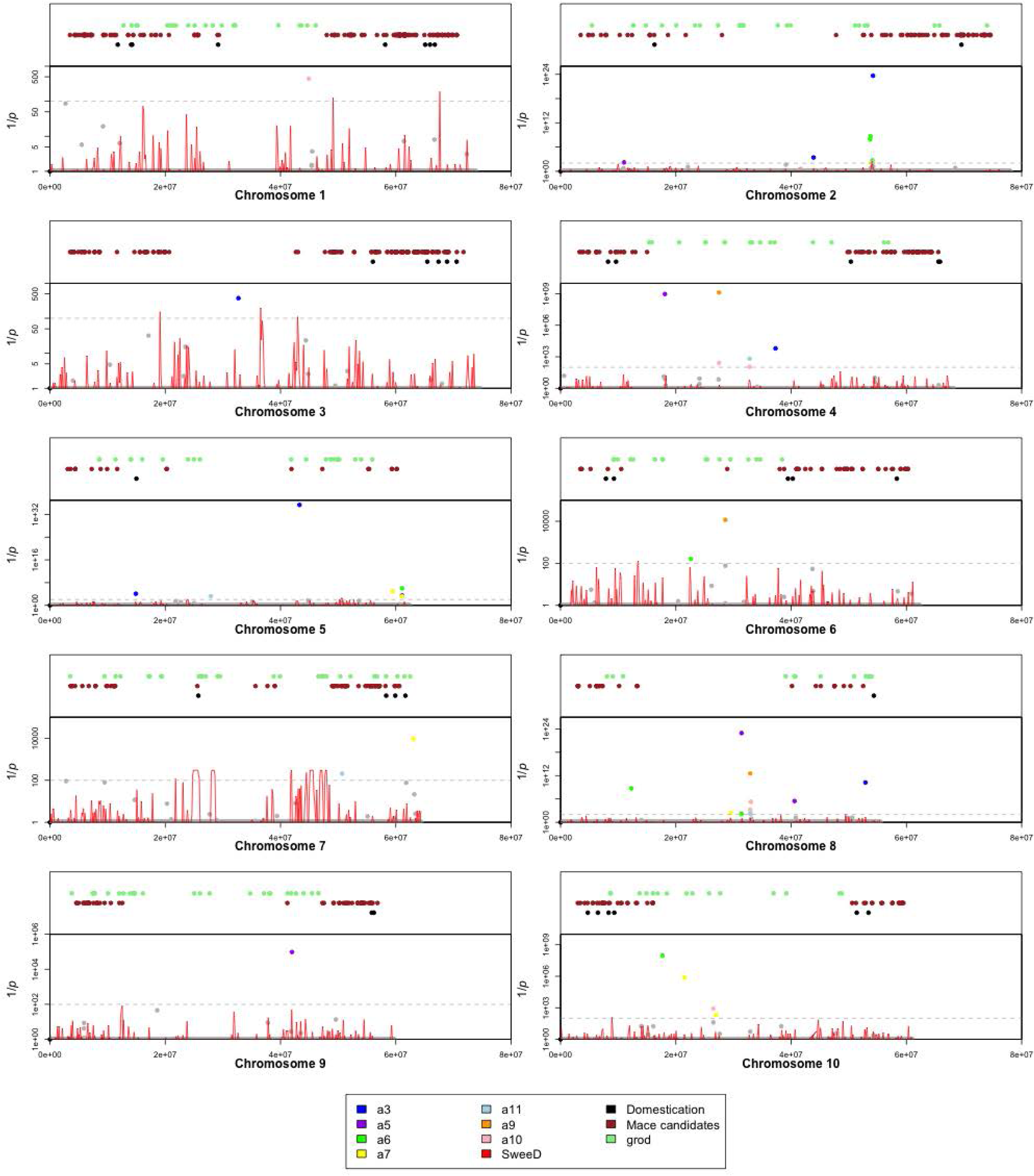
Selection signals across *S. bicolor* chromosomes 1 to 10. Heterozygosity ratio (wild/cultivated) inverted probabilities (Bonferroni corrected) shown in colours as described in key. Grey dashed line indicates 1% significance threshold after Bonferroni correction. SweeD values shown in red. Above: Locations of 38 known domestication genes shown in black. Locations of candidate domestication loci identified by Mace *et al* (24) shown in brown. Locations of GERP score regions of difference (grod) between genomes shown in green.

Together, the selection identification approaches exploit a range of different types of signature left by selection, and reveal a complex and dynamic history of selection over time summarized in Figure 5. We generally found more evidence for selection in the bicolor type sorghum than the durra type. Despite its apparent wild phenotype, the wild sorghum (A3) at Qasr Ibrim from 1765 yrs BP shows evidence of selection at domestication loci concerned with architecture (*int1, tb1*), suggesting possible introgression with contemporaneous domesticated forms (represented by sample A5) that could have contributed to reduced heterozygosity. Interestingly, the intensification signals show some overlap between samples A3 and A5 (*int1* and *ae1*), with A5 showing further evidence for selection at shattering, dwarfing and sugar metabolism loci (*Sh3/Bt1, dw2* and *SPS5*) that would contribute to the domesticated phenotype of A5 relative to A3. Subsequent to this a period of intensification in selection is apparent both for dwarfing and sugar metabolism traits (710-715 years BP) in the bicolor type, with ten domestication loci showing significantly low levels of heterozygosity in this lineage by 710 yrs BP in A7. In the bicolor type, two of the sugar metabolism associated gene families show evidence of early selection controlling photosynthetic sucrose production first (*SPS*) and then an intensification of selection for breakdown (*SUS*). A third gene family (*SUT*) associated with sucrose transport appears to come under later selection in bicolor. In contrast, fewer domestication loci were found to show evidence of intensifying selection in the durra type, and none showed evidence of low heterozygosity. In this case we detected signals for an intensification of selection on tillering and maturity associated loci (*gt1, ma3*, the latter also being detected using SweeD s1 in the bicolor lineage). Significant heterozygosity reduction was identified in windows containing a large number of disease resistance loci (pk4, pk11, pk15, pk20, pk24, pk25) as well as sugar metabolism loci (pk14, pk18, pk19, pk22) in the bicolor type. One of SweeD peak (s2) was closely matched to pk5 on chromosome 2 in the 54.0 – 54.2 Mbp interval, possibly indicating signatures for the same selection process. This region shows a consistently low heterozygosity over time in the bicolor type with the notable exception of the 1805 year old sorghum (A5). The region contains the far-red impaired response genes (*FAR1*), as well as anther indehiscence 1 (*AI1*). The *FAR1* gene is associated with phytochrome A signal transduction (22), so is important in responses to far red light that divert resources away from tall growth to increase root and grain growth. The *AI1* gene regulates anther development (23), allowing earlier development. Either of these genes may be locally adapted to the Qasr Ibrim environment since they already appear to be under intense selection in the wild sorghum at this site (sample A3), but not apparently under as much constraint in modern sorghum type bicolor.

The dynamic selection over time detected with most intensification of selection occurring before 1805 years BP, appears to correlate with a sharp increase in mutation load in the bicolor type. In contrast, the durra type shows much less evidence of selection and on arrival at Qasr Ibrim shows initially similar levels of mutation load to the bicolor type that then decreased over time (Figure S3). To investigate whether loci of selection are associated with higher regions of mutation load we measured the maximum deviations between genomes in GERP load scores across the genome and compared those to the locations of selection peak candidates (Figure 3, Table S9). Selection signatures were highly significantly associated with regions of maximum deviation in mutation load with 30% of low heterozygosity peaks (*p* 8.04 ×10^−9^), 45% SweeD peaks (*p* 2.55×10^−7^) and 26% domestication loci (*p* 5.03 × 10^−5^) occurring in such regions. The intensification of selection is associated with increased mutation load and could explain the spike in mutation load observed in the 1805 year old sorghum (sample A5).

### Genome rescue through hybridization

We considered that the decreasing mutation load observed in the durra type could be due to a genomic rescue caused by hybridization with the local bicolor type. To investigate for evidence of hybridization we first constructed a maximum likelihood phylogenetic tree of wild and cultivated total genomes (Figure S8), and individual trees for 970 sections across the genome (Supplementary data set 1). After accounting for biases introduced by ancient DNA modification, both the durra and bicolor type from Qasr Ibrim form a single clade to the exclusion of modern bicolor and durra types, suggesting they have indeed hybridized over time. D-statistic analysis for introgression (24) shows over time the durra type became increasingly similar to the local bicolor type, suggesting progressive introgression between the two types (Figure S9). We then compared the archaeological genomes against a global sorghum diversity panel (25,26) (Figure 4). The archaeological genomes are distributed along an axis of spread that has Asian durra types at the extremity. The oldest archaeological durra type (A11) sits between East African durra types and Asian durra types, whilst the wild phenotype sorghum, most closely aligned to the subsequent type bicolor, sits close to the center of the PCA, suggesting East African durras may have arisen from a hybridization between Asian durra and African bicolor. The oldest archaeological durra type in this study (sample A11) may represent one of the earliest of the east African durras. The younger archaeological genomes of the two types become progressively closer on the PCA supporting a process of ongoing hybridization between the two types over time.

**Figure 4.**
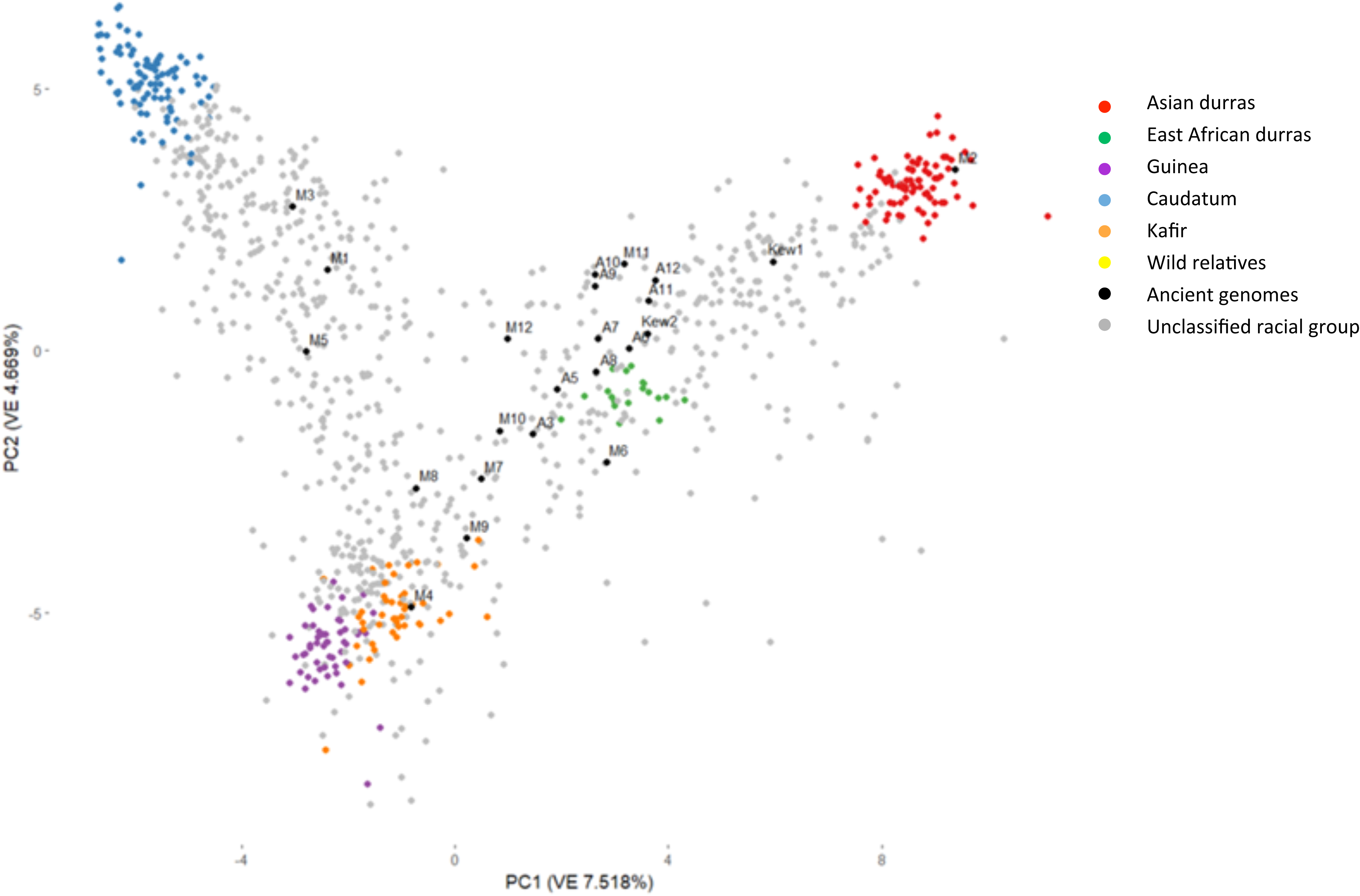
Principal Coordinate Analysis of 1894 SNPs from 23 genomes in this study and 1046 sorghum lines described in Thurber et al (25). Arrows indicate temporal movement of bicolor and durra type archaeogenomes in PCA.

**Figure 5.**
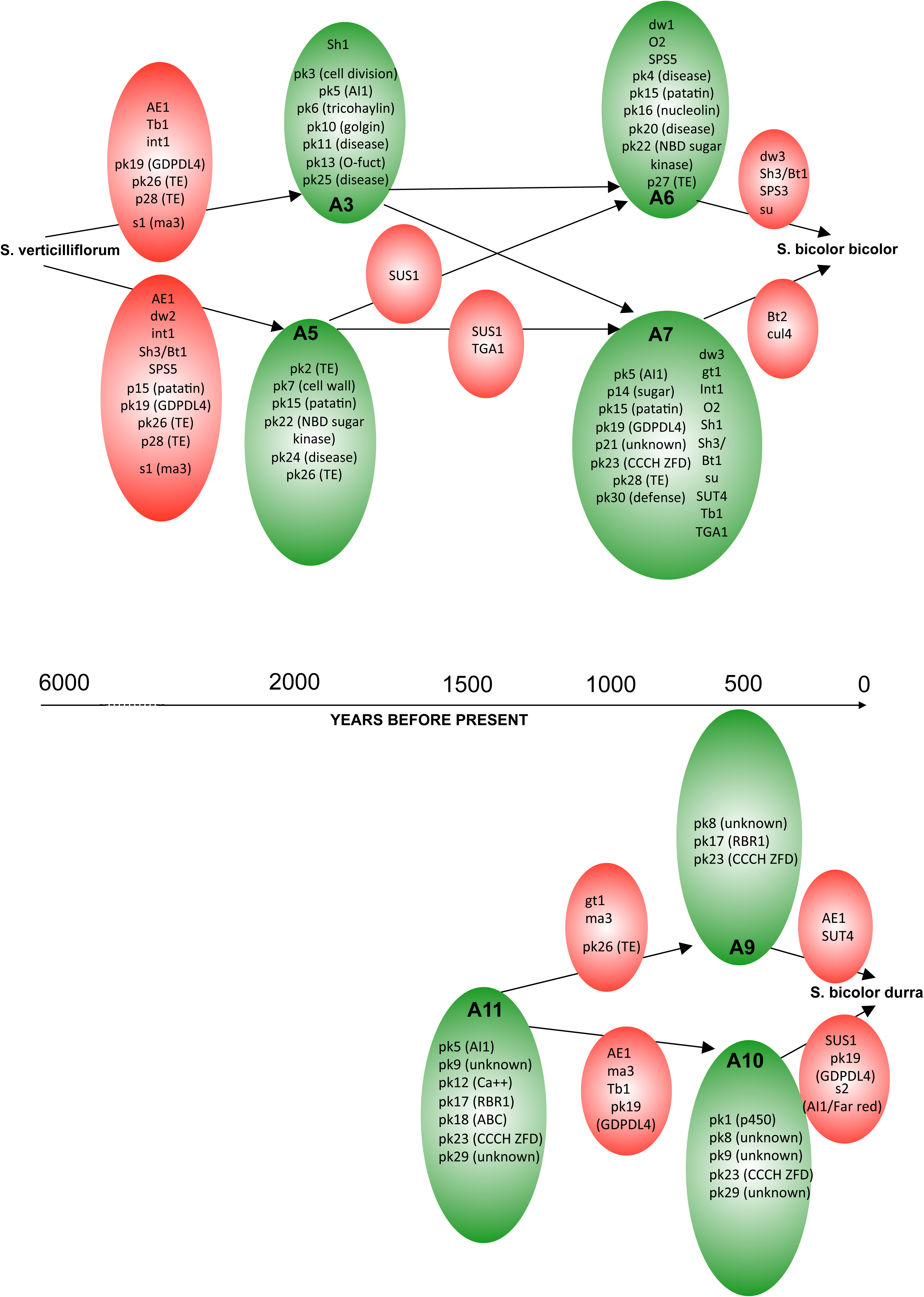
Summary of selection signals over time in archaeogenomes. Red indicates selection intensification episodes, green indicates selection signals identified by low heterozygosity or SweeD analysis.

Finally, we investigated whether the hybridization between the bicolor and durra types led to adaptive introgression or genomic rescue. Phylogenetic incongruence between the bicolor and durra type clades suggests that hybridization was frequent at loci under selection (Table S10). In agreement with previous studies (10) there is clear evidence for a donation of the dwarfing *dw1* allele from durra to bicolor with a single durra type sample sitting within the bicolor type clade in this region, but in most cases although the clades of bicolor and durra have become mixed, it is not sufficiently clear which is the more likely donor. Interestingly, seven of the nine sugar-metabolism associated loci potentially under selection in the bicolor type are also areas of introgression with durra. In all cases where identified was possible (*su, SUS1* and *SPS3*), durra was identified as the donor. However, in the case of *SPS5* in which we identified early intensification of selection in the bicolor type, no phylogenetic incongruence occurred. Conversely, at the maturity locus *ma3* containing region, the durra type A11 that was identified as potentially under selection sits within the bicolor clade suggesting a donation from bicolor to durra. The *FAR1/AI1* loci region, which appears to have been under strong selection in bicolor throughout, appears to have been donated from bicolor to durra.

Assuming that prior to the introduction of the durra type to Qasr Ibrim the two types, bicolor and durra, had accrued mutation loads independently, then hybridization would have afforded the opportunity for genomic rescue between the two types. We therefore considered all ancestor/descendent pairs of genomes within the bicolor and durra type lineages in the context of a third potential donor genome, and scanned all sites for comparative GERP load scores under the additive model. We calculated firstly the difference in GERP load scores between the ancestor and potential donor to give a ‘total rescue value’ that reflects a donor’s potential to reduce mutation load across the entire genome, Figure 5. We secondly assessed the donor’s potential to effect mutation load reduction specifically at only those sites in which there had been a reduction in GERP load score between the ancestor and descendent to give an ‘on target rescue value’.

In the case of durra sample A11 (1470 yrs BP) as ancestor and A9 as descendent, bicolor samples A6 (715 yrs BP) and A7 (710 yrs BP) are intermediate in age and therefore potential donor genome types from the bicolor lineage. The analysis predicts that either A6 or A7 would have reduced load in the regions that were observed to be reduced in the descendent A9 (505 yrs BP), but an overall detrimental effect to genome wide load, which is in fact observed (Figure S3). However, had earlier bicolor types been available for introgression, such as A5 (1805 yrs BP), then hybridization would have been more beneficial for the durra types. Generally, there is strong rescue potential of bicolor types by durra on the sites that were observed to improve, however in most cases there is an expectation that the over all load would be increased from the transfer of durra specific load to bicolor. Notably, durra types in general are predicted to reduce the on target load and genome wide load in A7, which is observed (Figure S3).

## Discussion

This study demonstrates that sorghum represents an alternative domestication history narrative in which the effects of a domestication bottleneck are not apparent, mutation load has accrued over time probably as a consequence of dynamic selection pressures rather than a domestication-associated collapse of diversity, and that genomic rescue from load occurred when two different agroclimatic types met.

The linear nature of the decreasing trend in diversity over time observed in sorghum in this study is surprising. An extreme bottleneck early in the history of would be expected to lead to a negative exponential trend as diversity is rapidly lost in the early stages of domestication. An alternative explanation for the trend could be that diversity has been lost steadily through drift over time. However, a simple drift model shows that such a ten-fold loss in diversity would also be associated with a negative exponential trend, Figure S11. It is possible that diversity loss could have been supplemented by gains through introgression from the wild over time, counteracting the trend made by drift. Sample A3 could be the result of a wild introgression event since there are older domesticate phenotypes in the archaeobotanical record, such as sample A5. Sorghum is known for its extensive introgression leading to a strong regional structure within cultivars (10), making continuous introgression seem like a plausible scenario for sorghum at Qasr Ibrim. Incorporation of three systems of introgression into the simple drift model in which introgression is either constant, diminishing or increasing over time still results in a non-linear trend, which become parabolic when introgression becomes very high over time (not shown), Figure S11. We therefore think it unlikely that a model of constant drift and introgression is causative of the apparent linear decrease in diversity over time observed in this study.

Such linear decreases in diversity have been observed in human populations with increasing geographic distance from Africa and are most robustly explained by sequential founder models (27). The annual cycle of crop sowing and harvesting also represents a serial founding event scenario. A simple model of founding events in which 25% of the harvest is set aside for sowing based on field experiments (28) demonstrates that loss of genetic diversity approximates a linear process as populations become large, Figure S12A, and that the gradient of diversity loss is highly correlated with the populations size, Figure S12B. On the basis of the gradient of diversity loss observed in sorghum, this model predicts a long-term population size of 289,407 for the sorghum in this study. This estimate is in excess of the effective population size estimated from the heterozygosity of wild sorghum, 135,823. It therefore seems plausible that in the case of sorghum diversity has likely been lost through a series of sequential founding episodes based on the cropping regime in a process that likely incorporated all the available wild genetic diversity at the outset rather than a substantial initial domestication bottleneck.

The deleterious effects of mutation load are becoming increasingly apparent and a major problem in modern crops such as the dysregulation of expression in maize (29). The study here demonstrates the potential immediacy of the problem in that mutation load may generally be a consequence of recent selection pressures leading to an exponentially rising trend rather than a legacy of the domestication process. While the general trend of the archaeogenomes is for the increase in the number of sites homozygous for deleterious variants (recessive model), the overall trend for the number of sites holding deleterious variants decreases (dominant model), which suggests a process of general purging of variants from the standing variation of the wild progenitor combined with the rise of homozygosity with decreasing diversity of the variant sites that remain. However, this is sharply contrasted by modern sorghum in which there is a leap in the number of sites holding deleterious mutations (dominant model). This process contributes to the accompanying jump in load under both the recessive and additive models in modern sorghum. This indicates a large influx of new deleterious variants within the last century giving the trend of mutation load accumulation an exponential shape. It is likely that this influx of mutation load is the product of recent breeding programs and the genetic bottlenecks associated with the Green Revolution. The accumulation of load has previously been associated with mutation meltdown and extinction of past populations (30) but it remains unclear whether crops could follow the same fate in the absence of rescue processes, or whether such episodes could have been involved with previous agricultural collapses when crops experienced extensive adaptive challenges (31,32). In the case of sorghum wild genetic resources may be valuable not only as a source of improved and environmentally adaptive traits, but also as a source for reparation of genome wide mutation load that may affect housekeeping and economic traits alike.

This represents the first plant archaeogenomic study that tracks multiple genomes to gain insight into changes in diversity over time directly. The trends revealed, based on a relatively low number of archaeological genomes, suggest a domestication history contrary to that typically expected for a cereal crop. Further archaeogenomes may establish whether this is a general trend for sorghum and other crops.

## Methods

### 1. Sample Acquisition

Archaeological samples were sourced from A. Clapham from the archaeological site Qasr Ibrim, outlined in Table S1. For details on dating see section 1.3 below. Historical samples from the Snowden collection were sourced from Kew Gardens, Kew1: Tsang Wai Fak, collection no. 16366 Kew2: Tenayac, Mexico, collection assignation ‘s.n.’. Modern samples of *S. bicolor* ssp. *bicolor* type bicolor, durra, kafir, caudatum, drumondii and guinea were supplied through the USDA [accession numbers PI659985, PI562734, PI655976, PI509071, PI653734 and PI562938 respectively]. Wild sorghum samples *S. vertilliciliflorum, S. arundinaeum, and S. aethiopicum* were also obtained from the USDA [accession numbers PI520777, PI532564, PI535995], and wild *S. virgatum* was donated by D. Fuller. The outgroups *S. propinquum* and *S. halapense* were obtained from the USDA [accession numbers PI653737 and Grif 16307] respectively.

The genomes generated in this study were also compared to 1023 re-sequenced genomes taken from Thurber et al 2013 (26).

#### 1.2 A note on taxonomy

The sorghum genus is complex with numerous taxonomic systems. After Morris *et al.*’s findings (10), we have elected not to describe the principal cultivar types as subspecies or races but rather simply ‘types’ to reflect the reality that there is evidence of considerable introgression between each of these forms. The wild progenitor of domesticated sorghum is a complex made up of four ‘races’ verticilliflorum, arundinaceum, aethiopicum and virgatum. However, the integrity of these races is also questioned, and the currently more accepted designation is one species, verticilliflorum, of which the other races are subtypes. For clarity and simplicity in this study we have used the race type as a variety designation.

#### 1.3 A note on Qasr Ibrim and archaeological context of samples

Qasr Ibrim was a fortified hilltop site in the desert of Lower Nubia on the east bank of the Nile, about 200 km, south of Aswan in modern Egypt. It has been excavated over numerous field seasons, since 1963 by the Egyptian Exploration Society (UK). In recent years with higher Lake Nasser levels only upper parts of the site are preserved as an island (33,34). The desert conditions provided exceptional organic preservation by desiccation with exceptional preservation of a wide range of biomolecules (e.g. 35-37). Systematic sampling for plant remains was initiated in 1984 (38) and the first studies of these remains were carried out in the 1980s by Rowley-Conwy (39) and had continued by Alan Clapham (40,41). The exceptional plant preservation has previously allowed successful ancient genomic studies of barley (35) and cotton (36).

Qasr Ibrim was founded sometime before 3000 years BP. It had occupations associated the Napatan kings (Egyptian Dynasty 25: 747-656 BC), possible Hellenistic and Roman Egypt (3^rd^ century BC to 1^st^ c. AD), the Meroitic Kingdom (1^st^ century to 4^th^ century AD), and local post-Meroitic (AD 350-550) and Nubian Christian Kingdoms (AD 550-1300). Earlier periods are associated temples to Egyptian and Meroitic deities. After Christianity was introduced the site had a Cathedral. Later Islamic occupations finished with use as an Ottoman fortress. The site was abandoned in AD1812. The Sorghum material studied here comes from a range of different contexts from excavation seasons between 1984 and 2000. While the chronology of the site is well established by artefactual material, including texts in various scripts, several sorghum remains or associated crops, were submitted for direct AMS radiocarbon dating, as listed below in Table S2. For directly dated find the median of the 2-sigma calibrated age range has been used. Note that Radiocarbon calibration defines “the present” as AD 1950, and we have recalculated the median as before AD 2000, and assigned Snowden historical collections form the start of the 20^th^ century as ca. 100 BP. For material not directly dated, sample A12 could be assigned based on associated pottery and finds, which have a well-established chronology through the Christian periods (42), A12 is associated with Islamic/Ottoman material (1500-1800 AD, ca. 400 BP)

### 2. DNA extraction

DNA was extracted from archaeological and historical samples in a dedicated ancient DNA facility physically isolated from other laboratories. All standard clean-lab procedures for working with ancient DNA were followed. Single seeds from each accession were ground to powder using a pestle & mortar and incubated in CTAB buffer (2% CTAB, 1%PVP, 0.1M Tris-HCl pH 8, 20mM EDTA, 1.4M NaCl) for 5 days at 37°C. The supernatant was then extracted once with an equal volume of 24:1 chloroform:isoamyl alcohol. DNA was then purified using a Qiagen plant Mini Kit with the following modifications: a) 5x binding buffer was used instead of 1.5x and incubated at room temperature for 2 hours before proceeding. b) After washing with AW2, columns were washed once with acetone and air-dried in a fume hood to prevent excessive G-forces associated with centrifugal drying. c) DNA was eluted twice in a total of 100µl elution buffer and quantified using a Qubit high sensitivity assay.

DNA from modern samples was extracted using a CTAB precipitation method due to excessive polysaccharide levels precluding column-based extractions. Briefly, seeds were ground to powder and incubated at 60°C for 1 hour in 750 ul CTAB buffer as previously described, with the addition of 1ul β-mercaptoethanol. Debris was centrifuged down and the supernatant was extracted once with an equal volume of 24:1 chloroform:isoamyl alcohol. The supernatant was then collected and mixed with 2x volumes precipitation buffer (1% CTAB, 50mM Tris-HCl, 20nM EDTA) and incubated at 4°C for 1 hour. DNA was precipitated at 6°C by centrifugation at 14,000 *g* for 15 minutes. The pellet was washed once with precipitation buffer and incubated at room temperature for 15 minutes before being centrifuged again under the same conditions. The pellet was dried and resuspended in 100µl high-salt TE buffer (10mM Tris-HCl, 1M NaCl) and incubated at 60°C for 30 minutes with 0.5µl RNase A. The DNA was then purified using Ampure XP SPRI beads.

### 3. Library construction and genome sequencing

Libraries for all samples were constructed using an Illumina TruSeq Nano kit, according to manufacturers’ protocol. A uracil-intolerant polymerase (Phusion) was used to amplify the libraries, in order to eliminate the C to U deamination signal often observed in ancient DNA in favour of the 5’ 5mC to T deamination signal. The purpose of this was to obtain epigenomic information after analysis using epiPaleomix (43). Consequently the data set was reduced for non-methylated cytosine deamination signals in the 5’ end, but showed expected levels of G to A mismatches for ancient DNA (5-10%) in the 3’ end and high levels of endogenous DNA content typical for samples from this site (Table S1). While this approach is thought to reduce library complexity by reducing the number of successfully amplified molecules, we considered this to be a worthwhile trade-off considering the exceptional preservation and endogenous DNA content of the Qasr Ibrim samples. We found no evidence to suggest insufficient library complexity after amplification. A minor modification was made to the protocol for ancient and historical samples: a column-based cleanup after end repair was used, in order to retain small fragments that would otherwise be lost under SPRI purifications as per the standard protocol. Genomes were sequenced on the Illumina HiSeq 2500 platform. Ancient and historical samples were sequenced on one lane each using SR100 chemistry and modern samples on 0.5 lanes each using PE100 chemistry.

### 4. Preliminary Bioinformatics processing

Illumina adapters were trimmed using cutadapt v1.11 using 10% mismatch parameters. Resulting FastQ files were mapped to the BTX623 genome (44) using bowtie2 v2.2.9 (46) under--sensitive parameters. SAM files containing mapped reads with a minimum mapping score of 20 were then converted to BAM files using samtools v1.14 (47). Variant calls format (VCF) files were then made from pileups constructed using samtools mpileup, and variant calls were made using bcftools v1.4 (47).

### 5. Methylation analysis

Since a uracil-intolerant polymerase was used for library generation, we analysed BAM files using epipaleomix (43) on the ancient samples. We then collated the number of identifiable 5mC sites globally for each sample. Epipaleomix is designed to characterise CpG islands typical to animal genomes and, is not suited to gene-specific analysis of plant genomes to due to their wider methylation states (CHH and CHG) (45). However when assessing relative overall genome methylation between individuals of the same species, CpG islands measured in this way provide a perfectly adequate proxy. We opted for global and windowed-measurements to determine relative methylation states between samples.

### 6. Evolutionary and population analyses

Two archaeological genomes (A8 and A12) were from phenotypes intermediate between bicolor and durra types. We found that sample A8 was predominantly of bicolor type and A12 predominantly of durra type. Given the uncertainty of these samples and their likely hybrid origins, we elected to leave them out of most analyses.

#### 6.1 Heterozygosity analysis

The number of heterozygous sites was measured for each 100 kbp window of genome aligned to the BTX_623 reference sequence (44). The frequency distribution of heterozygosity was then calculated by binning the windows in 1 heterozygous base site intervals. Ratios of wild:cultivated heterozygosity were calculated for each window using *S. verticilliflorum* as the wild progenitor. Ratios closely approximate a negative exponential distribution. Probabilities of observed heterozygosity ratios for each window were obtained from a negative exponential distribution with λ equal to 1/µ for all ratios for each chromosome. A Bonferroni correction was applied by multiplying probability values by the number of windows on a chromosome in Figure 4. Locations of 38 known domestication syndrome loci (shown in Tables S5 and S7) were obtained by reference to the BTX_623 genome. Candidate domestication loci were obtained from the scans of Mace *et al* (25). In the genome-wide scan peaks were considered significant if 1/*p* > 100 after Bonferroni correction.

We considered the possibility that the observed heterozygosity levels may be influenced by postmortem DNA damage. To explore this, we characterized the relationships between time, heterozygosity and postmortem deamination. As we previously described, C to U damage signals are eliminated at the 5’ ends of sequence reads because of our choice of polymerase, so we therefore characterized damage profiles at the 3’ ends only, using mapDamage output statistic ‘3pGtoA_freq’ and taking a mean of the 25 reported positions for each ancient or historical sample. Unsurprisingly, we found that the accumulation of damage patterns is a function of time in a logistic growth model, assuming a zero-point intercept for both factors (R^2^ = 0.9). 80% of damage capacity under this model is reached reasonable quickly, in 331.0 years. All the Qasr Ibrim samples are at least 400 years old, and so we re-fitted a linear regression model to these samples only so characterize these relationships in a true time-series. We found a negligible correlation between time and damage accumulation after 400 years (R^2^ = 0.15, p = 0.34). Next, we characterized the relationship between age and heterozygosity under the same model (although without the assumption of a zero-point intercept, since even modern domesticate lines in this study show non-zero levels) and found a weak fit (R^2^ = 0.64, p = 0.14). This relationship is however likely influenced by our central hypothesis, with ‘less domesticated’ samples being earlier in the archaeological record, and so a counter-argument should not be inferred from this analysis. Finally, we assessed the relationship between damage and heterozygosity by linear regression, assuming inappropriateness of a logistic model since both damage and heterozygosity factors are functions of time. We found a weak correlation when considering all samples (R^2^ = 0.2, p = 0.2), and virtually no correlation when considering the Qasr Ibrim time series only (R^2^ = 0.04, p = 0.61). Considering that the two historical Kew samples are ostensibly domesticates, and historical and geographic outliers to the rest of the dataset, we conclude that the observed levels of heterozygosity in the ancient samples are not influenced by postmortem damage patterns.

#### 6.2 Differential Temporal Heterozygosity Gradient Analysis

Our rationale was to utilize the temporal sequence of genomes to identify time intervals associated with intensification of selection. To this end we designed an analysis to identify outliers in changing heterozygosity over time to the general genomic trend. We considered all possible historical paths between genomes given three pairs of samples were almost contemporaneous (A3/A5, A6/A7 and A9/A10), with wild *S. verticilliflorum* representative of the wild progenitor in the case of the bicolor lineages.

For each 100kbp window we calculated the gradient of change in heterozygosity between temporally sequential genome pairs by subtracting younger heterozygosity values from older and dividing through by the time interval between samples. Genome-wide gradient values for all 100kbp windows were used to construct a non-parametric distribution to obtain probability values of change over each time interval for a 100kbp window between a particular pair of samples. Peak regions identified by heterozygosity ratio, SweeD analysis and known domestication syndrome genes were then measured for gradient probability.

#### 6.3 SweeD analysis

VCF files from our 23 ancient, historical and modern samples and also 9 samples from Mace et al (25) were combined using the GATK (52) program CombineVariants. Subsequently, the combined VCF file was filtered - using bcftools v1.4 (47) - to only include sites with 2 or more distinct alleles and at sites where samples have depth less than 5 or a variant calling quality score less than 20 to exclude those samples. Then a further filter was applied - using bcftools v1.4 - to exclude variant calls due to C->T and G->A transitions relative to the reference, which potentially represent post-mortem deamination which has a high rate in aDNA samples (48). SweeD (21) was run with options for multi-threading (to run with 64 threads) and to compute the likelihood on a grid with 500 positions for each chromosome.

#### 6.4 Genome Evolutionary Rate Profiling (GERP) analysis

This analysis was carried out broadly following the methodology of Cooper *et al.* (19). We aligned the repeat-masked genomes of 27 plant taxa to the BTX_623 sorghum reference genome using last, and processed resulting maf files to form netted pairwise alignment fastas using kentUtils modules maf-convert, axtChain, chainPreNet, chainNet, netToAxt, axtToMaf, mafSplit, and maf2fasta. We forced all alignments into the frame of the sorghum reference using an expedient perl script, and built a 27-way fasta alignment excluding sorghum for GERP estimation. We created a fasta file of fourfold degenerate sites from chromosome 1 (347394 sites; NC_012870) with a perl script, and calculated a neutral rate model using phyloFit, assuming the HKY85 substitution model and the following tree:

*(((((((((Trifolium_pratense,Medicago_truncatula),Glycine_max),Prunus_persica),(Populus_trichocarpa,Manihot_esculenta)),(((Arabidopsis_thaliana,Arabidopsis_ly rata),(Brassica_napus,Brassica_rapa)),Theobroma_cacao)),Vitis_vinifera),((Sola num_tuberosum,Solanum_lycopersicum),(Chenopodium_quinoa,Beta_vulgaris))), (((Zea_mays,Setaria_italica),(((Oryza_rufipogon,Oryza_longistaminata),Leersia_ perrieri),(((Triticum_urartu,Aegilops_tauschii),Hordeum_vulgare),Brachypodium_ distachyon))),Musa_acuminata)),Amborella_trichopoda)*

We then calculated GERP rejected subsitutions (RS) scores using gerpcol with the default minimum three taxa represented for estimation. The mutation load for each genome was then assessed by scanning through their VCF files generated by alignment to BTX_623. Maize was used as an outgroup to judge the ancestral state, and only sites at which there was information from maize were incorporated into the analysis. Sites which differed to the ancestral state were scored based on the associated RS score for that site following the scheme of Wang *et al.* (18): 0, neutral, 0-2 slightly deleterious, 2-4, moderately deleterious, >4 seriously deleterious. We collected scores under three models, recessive, additive and dominant. Under the dominant model we counted each site once regardless of whether it had one or two alternative bases to the ancestor, so giving the total number of base sites containing at least one potentially deleterious allele. Under the additive model we counted the total number of alleles that were alternative to the ancestor such that each homozygous alternative site scored 2, but heterozygous sites scored 1. Under the recessive model only sites that were homozygous for potentially deleterious variants were counted.

To investigate the significance of overlap between regions significant GERP regions of difference (GROD) between taxa and signatures of selection we used a binomial test in which the null probability of selecting a GROD was equal to the total number of GRODS (193) divided by the total number of 100 kbp regions studied (6598), and N and *x* were the total number of selection signals and the number of selection signals occurring in a GROD respectively.

We used the GERP profiles to explore potential genomic rescue from mutation load accrued independently in the bicolor and durra lineages prior to hybridization between the two types. For the purposes of this analysis we used the wild sorghum genome A3 as a possible wild ancestor genome to the domesticated bicolor form A5 even though this wild sample is contemporaneous to that domesticated form. All possible ancestor descendent pairs were assembled within bicolor or durra types, and all 100 kbp windows were scanned for the relative additive model GERP load scores for ancestor, descendent and a third potential donor genome. The total potential for the donor genome to rescue the ancestral genome was scored summing the difference in GERP scores across all windows between the ancestor and donor. To better fit a scenario in which the donor genome was the causative agent of reduction GERP load score we identified windows that satisfied the condition ancestor GERP load score > descendent GERP load score, and summed up the difference in ancestor and potential donor scores to give an ‘on target rescue’ value.

#### 6.5 Phylogenetics

Maximum likelihood tress were constructed using exaML (49) firstly using whole genome sequences (Figure S7), and for 970 consecutive blocks across the genome (supplementary data set). Prior to computing phylogenetic trees, the VCF files were processed as described in section 6.3 (on the SweeD analysis) albeit with our 23 ancient, historical and modern samples only.

The maximum likelihood tree using the whole genome sequences was constructed as follows. Our own script created a multiple sequence alignment file by concatenating the variant calls in the VCF file and outputting the results in PHYLIP (50) format. The program parse-examl from the ExaML package (version 3.0.15) was run in order to convert the PHYLIP format file into ExaML’s own binary format. Also, ExaML requires an initial starting tree which was obtained by running (on multiple threads) Parsimonator v1.0.2, a program available as part of the RaxML package (51) - developed by the same research group - for computing maximum parsimony trees. An ExaML executable (compiled to run using MPI) was run on multiple CPUs in order to compute the maximum likelihood tree.

The trees for 970 consecutive blocks across the genome were computed by essentially the same approach as described above for a single tree, after a script obtained the blocks from the input VCF file (for the combined samples) and output them in PHYLIP format.

To assess potential donation between genomes at candidate loci we examined trees spanning the corresponding100kbp windows. The tree topology was examined for congruence in the maintenance of bicolor and durra type groups within the Qasr Ibrim group of genomes. Instances of phylogenetic incongruence were interpreted as candidate regions of recombination between the two genome types, although identification of the donor and recipient genomes was not always clear. Simple cases in which a single genome from one sorghum type was found within the group of the other type were interpreted as possible genome donations from that group to the single genome. In the case of regions that scored highly in the SweeD analysis no phylogenetic congruence was attempted because the taxon in which selection has operated is not identified.

#### 6.6 Principal Component Analysis of global diversity set

A subset of 1894 SNPs were used to find the principal axes of genetic variation for the 23 samples and an unpublished set of 1046 diverse sorghum lines spanning the racial and geographic diversity of the primary gene pool of cultivated sorghum. 580 of these diverse lines were described in Thurber et al (26). These lines were produced within the Sorghum Conversion Program which introgressed key height and phenology genes into exotic lines to enable them to be produced in sub-tropical environments. The introgressed regions spanned approximately 10% of the genome which were masked for the purposes of this analysis. Principal component analysis of the centered data matrix was performed in R (R core team, 2017) using the *prcomp* function in the base “stats” package.

#### 6.7 D statistics

Patterson’s D-Statistic and modified F-statistic on Genome wide SNP data was used to infer patterns of introgression (24). D-statistic and fd-statistic for each of the 10 chromosomes was calculated using the R-package PopGenome. Variant Call Format (VCF) file, which is generated after mapping reads of an individual sample to the reference genome, was given as input to the readVCF() function of the package (52).

We used four R-language based S4 class methods from PopGenome package to carry out the introgression tests for every chromosome. First, we used the method set.population by providing 3 populations (2 sister taxa and an archaic group) viz., P1=BTX_623, P2=varying samples, P3=Most ancient S.bicolor A3. Second, using set.outgroup function, we set an outgroup (P4= S.halapense). Third, the method introgression.stats was employed to calculate the introgession tests. Finally, we used jack.knife.transform method (53) which transforms an existing object belonging to GENOME class into another object of the GENOME class with regions that corresponding to a Jackknife window. Standard error was then calculated by eliminating one such window i.e., a single chromosome under study and calculation was applied to the union of all the other chromosomes.

We tested for admixture from the most ancient *S. bicolor* type bicolor sample (A3), assuming this represents a genome prior to the appearance of the durra type on the African continent. The BTX_623 sorghum reference genome was taken as P_1_, sample A3 was taken as P_3_ and *S. halapense* was taken as the out group P_4_. *S. halapense* is native to southern Eurasia to east India and does not readily cross with *S. bicolor.* Samples were then tested at the P_2_ position across all 100kbp windows, each chromosome tested separately. Negative values (indicating an excess of P_1_/P_3_ combinations) are expected when the BTX_623 genome is more similar to sample A3 than P_2_. This is observed as expected for the durra types, although the value of D decreases over time consistent with either an increase in instances of P_2_/P_3_ or instances of P_1_/P_2_, both suggesting progressive introgression between the durra and bicolor types over time. Positive values (indicating an excess of P_2_/P_3_ combinations) suggest a close relationship between sample A3 and P_2_, which is observed the Qasr Ibrim bicolor types (A5, A6 and A7).

#### 6.8 Linear and exponential line fitting to heterozygosity and GERP score data

A straight line was fit to the heterozygosity data in Figure 2 using the glm function (for generalized linear models) in R and also an exponential function was fit to the same data using the gnm package (for generalized non-linear models) in R obtaining the values for the parameters, standard errors, p and AIC shown in Table S3. (It was confirmed similar values were obtained for the parameters, standard errors, p and AIC by fitting the straight line model using the gnm package in R.)

#### 6.9 Basic simulation of diversity loss through drift, introgression and serial founding events

To explore the effect on general trend line shape of introgression over time we used a basic simulation of drift loss using the standard equation:

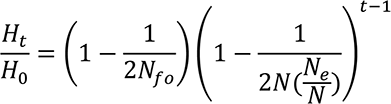

where *N*_*fo*_, *N* and *N*_*e*_ are the founding population size, census population size and effective population size respectively. For simplicity, we assumed in the case of our crop that all three population sizes were equal. To incorporate introgression we used a simulation to calculate and modify each generation by using the above equation to modify the diversity from the previous generation, and then adding a diversity value representative of gene flow. Gene flow was altered each generation by a power factor *f*, which was 1 in the case of constant introgression, 1.0001 in the case of diminishing introgression over time and 0.99995 in the case of increasing introgression over time, with an initial value for introgression as 0.000015, equating to the value of genetic diversity added to the population each generation. We used a founder population of 2000 for 6000 generations to recapitulate the observed 10-fold loss of diversity over this time frame in sorghum.

The serial founder event simulation was executed using the following model: To initialize, allele frequencies were randomly assigned for a defined number of alleles using a uniform distribution. We also applied a skewed distribution in which the first allele frequency generated as above was amplified to become a dominating allele frequency by a defined value. We found this made no difference to the simulation outcomes. N individuals were then randomly drawn using the allele frequencies, and the resultant frequency distribution calculated. Homozygosity was calculated as the sum of the squares of allele frequencies and subtracted from 1 for the heterozygosity. To convert these allele values to a per base site heterozygosity comparable to the sorghum data we divided by 1/He of the wild progenitor. A founder event was then generated by drawing Nb individuals from the allele frequency distribution, the new resultant allele frequency distribution calculated. N individuals were then drawn from this distribution, new frequencies calculated and the resultant heterozygosity calculated as above. The process was repeated for a defined number of cycles. We explored a scenario in which the founder population was based on setting aside 25% of the (seed) population each year, following classic experimental archaeology field trials (28). We explored several orders of magnitude of N (100, 1000, 10000, 100000), and assumed 5 alleles per gene, for 1000 founding events, equating to 1000 years of agriculture. Each trial was repeated 100 times, equating to 100 genes being simulated independently. While the overall distribution of diversity loss over time is exponential, seen more clearly at smaller population sizes, the trend approximates linear more closely with increasing population size (Figure 12A). We calculated the gradient of descent from the first 60 founding events and found the logs of the gradient and population size to be directly proportional (Figure 12B). We used linear regression of this relationship to predict the log of the population size associated with the log of the observed gradient of descent of diversity in sorghum. We independently calculated the effective population size associated with wild sorghum using the heterozygosity as an estimate of θ (using the relationship θ=4Neµ), and an estimate of 5 ×10^−9^ subs/site/year for the neutral mutation rate (54).

## Acknowledgements

The authors would like to thank M. Nesbitt for permitting the use of herbaria material from Kew. OS, WN, GB and RGA were supported by the NERC (NE/L006847/1) and LK was supported by NERC (NE/L012030/1). CJS and DQF work with archaeobotanical materials was supported by a European Research Council grant (no. 323842). Sequence data were deposited in the European Molecular Biology Laboratory European Bioinformatics Institute [project code PRJEB24962.].

---

Table S1. Sample details of genomes sequenced in this study

Table S2. Radiocarbon dates on sorghum specimens or closely associated plant remains

Table S3 Summary statistics of fit to linear and exponential models of change in heterozygosity and GERP score in *S. bicolor bicolor.*

Table S4. Peaks of significant heterozygosity ratios (wild/cultivated) and associated annotated gene models. * refers to genes also found in Mace et al (24) candidate domestication loci gene set.

Table S5. *P* values of heterozygosity ratio peaks.

Table S6 *P* values of heterozygosity ratio peaks for known domestication loci.

Table S7 Peaks of high signal found with SweeD analysis and associated annotated gene models. * refers to genes also found in Mace et al (24) candidate domestication loci gene set.

Table S8 *P* values of gradient deviation in heterozygosity over time relative to genomic average.

Table S9 Genomic locations for 100kbp windows that differ in gerp load between genomes by more than two standard deviations.

Table S10 Phylogenetic congruence between type bicolor and type durra clades in selection candidate regions.

**Figure S1.**
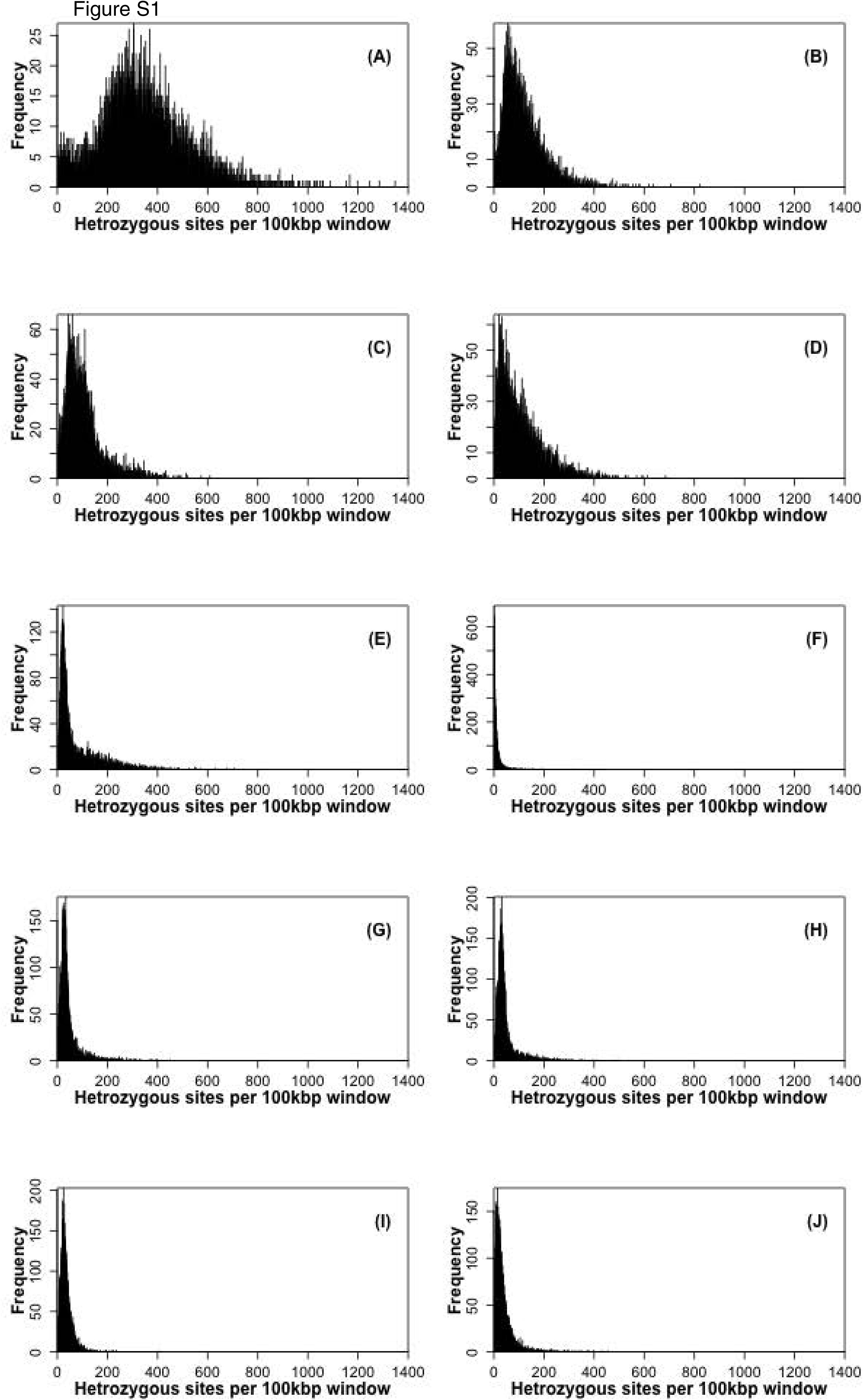
Frequency distributions of heterozygosity in genomes for 100 Kbp windows. A. *Sorghum verticilliflorum*. B. *S. bicolor* ‘wild phenotype’ (sample A3) 1765 yrs BP, C. *S. bicolor* type bicolor (sample A5) 1805 years BP, D. *S. bicolor* type bicolor (sample A6) 715 years BP, E. *S. bicolor* type bicolor (sample A7) 710 years BP, F. *S. bicolor* type bicolor BTX 623, G. *S. bicolor* type durra (sample A11) 1470 years BP, H *S. bicolor* type durra (sample A9) 505 years BP, I S. *bicolor* type durra (sample A10) 450 years BP, J *S bicolor* type durra modern.

**Figure S2.**
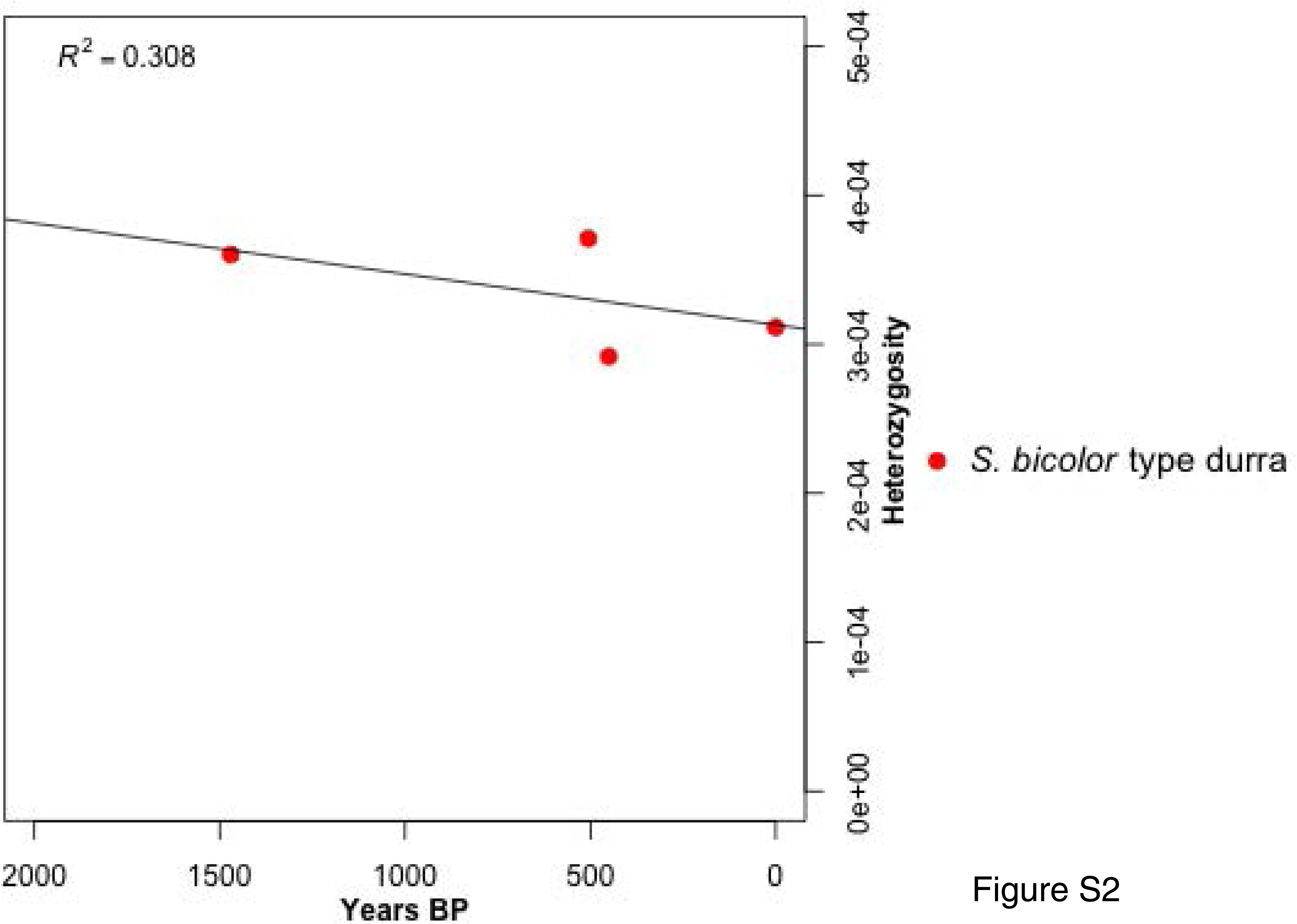
Heterozygosity over time in *S bicolor* type durra

**Figure S3.**
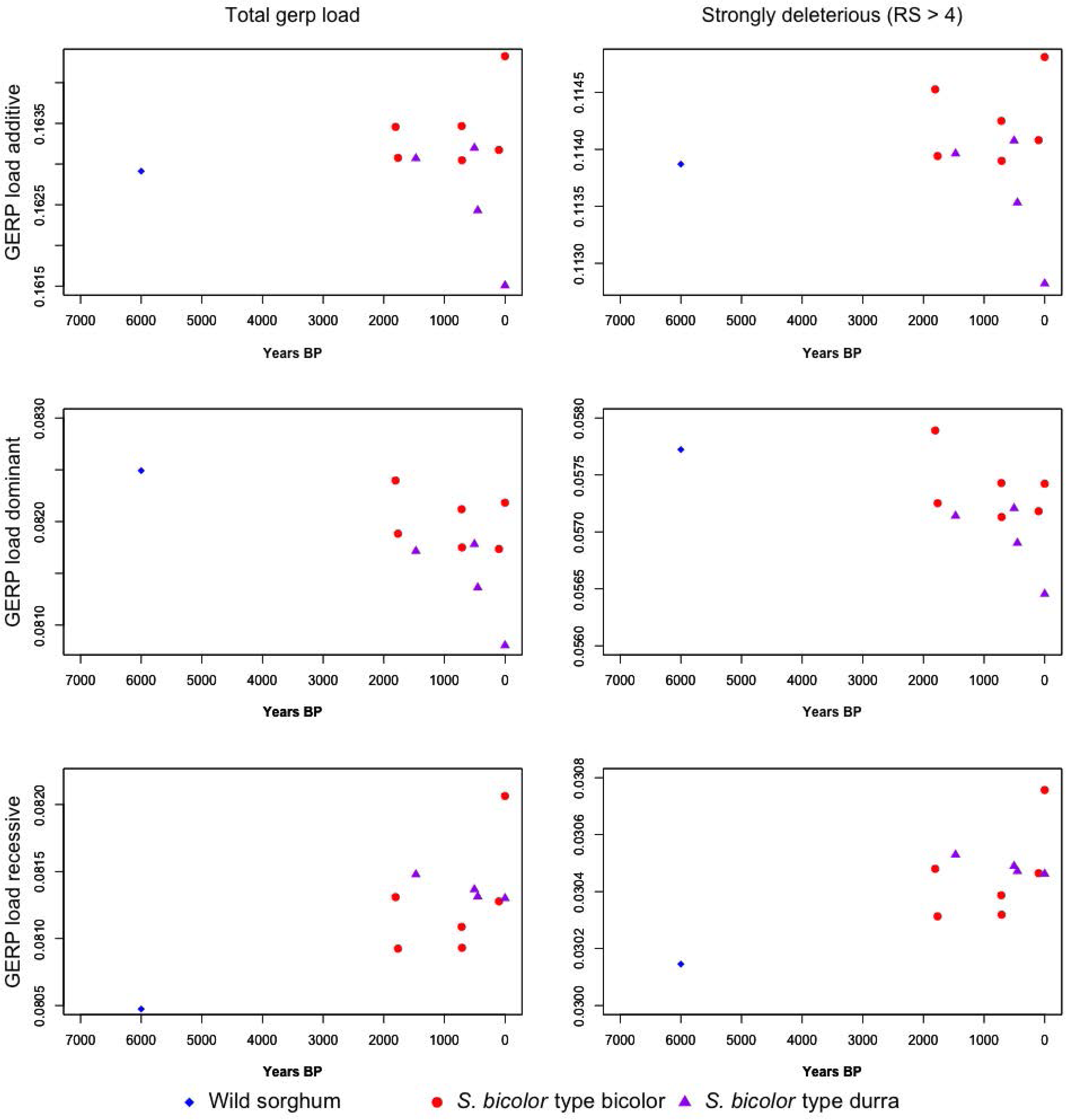
Additive, dominant and recessive model GERP load scores in *S bicolor* type bicolor and *S bicolor* type durra over time. Total GERP load calculated from variant sites with RS scores > 0, strongly deleterious GERP load calculated from variant sites with RS scores > 4. See methods for details on models.

**Figure S4.**
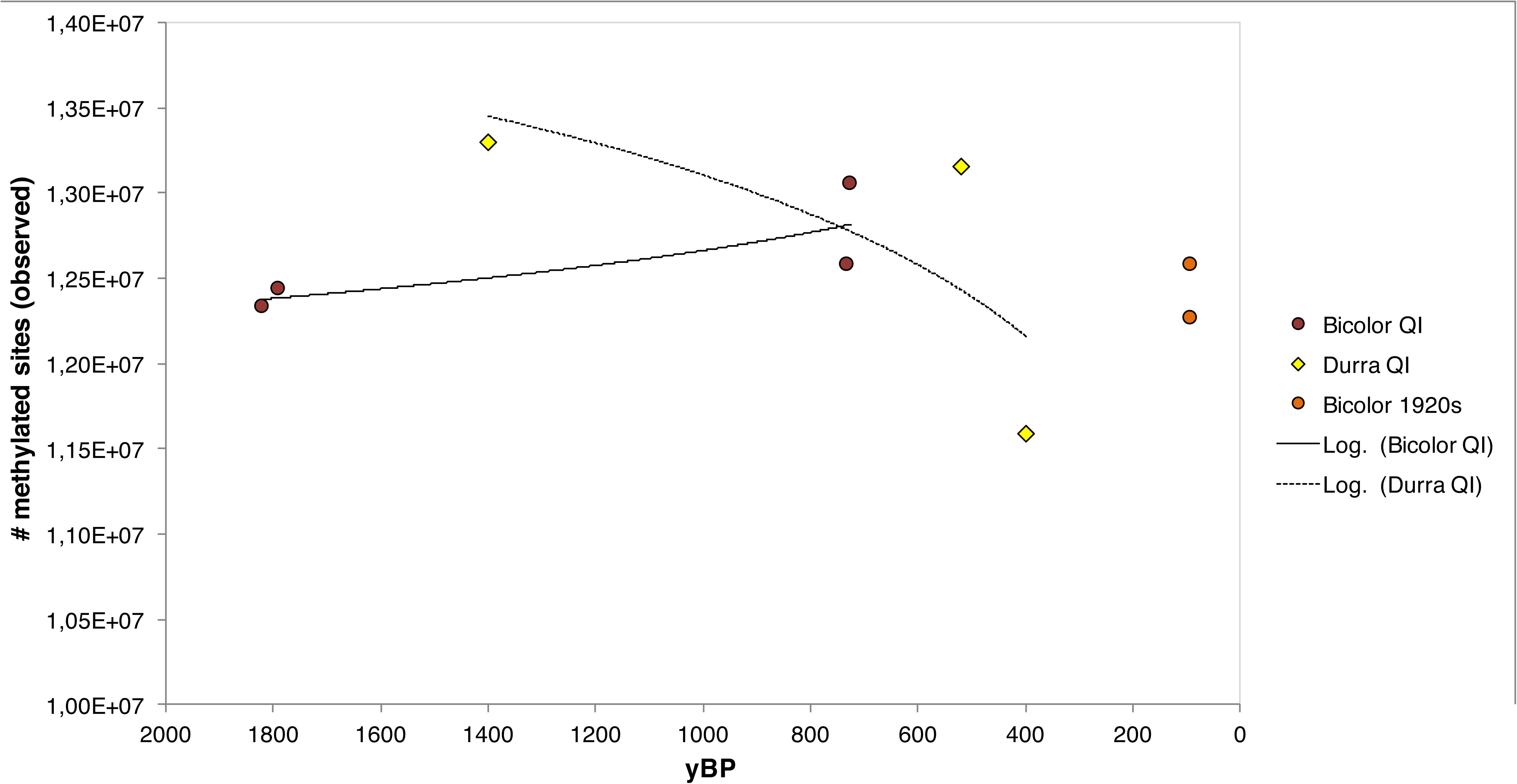
Methylated site number in *S bicolor* type bicolor and *S bicolor* type durra over time.

**Figure S5.**
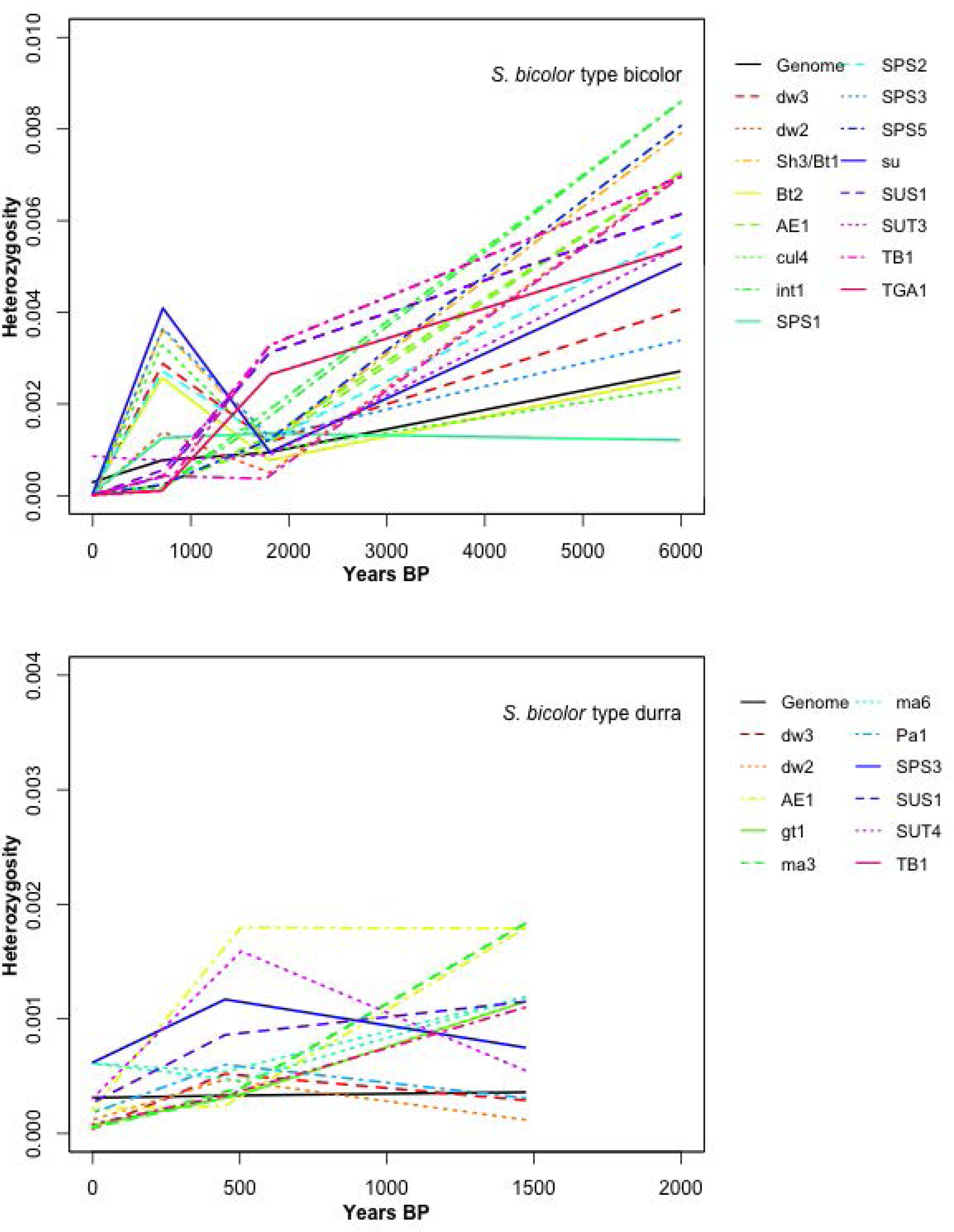
Heterozygosity over time of regions containing genome-wide significant wild/cultivated ratios. Significant deviations from the genomic gradient of change over time shown only.

**Figure S6.**
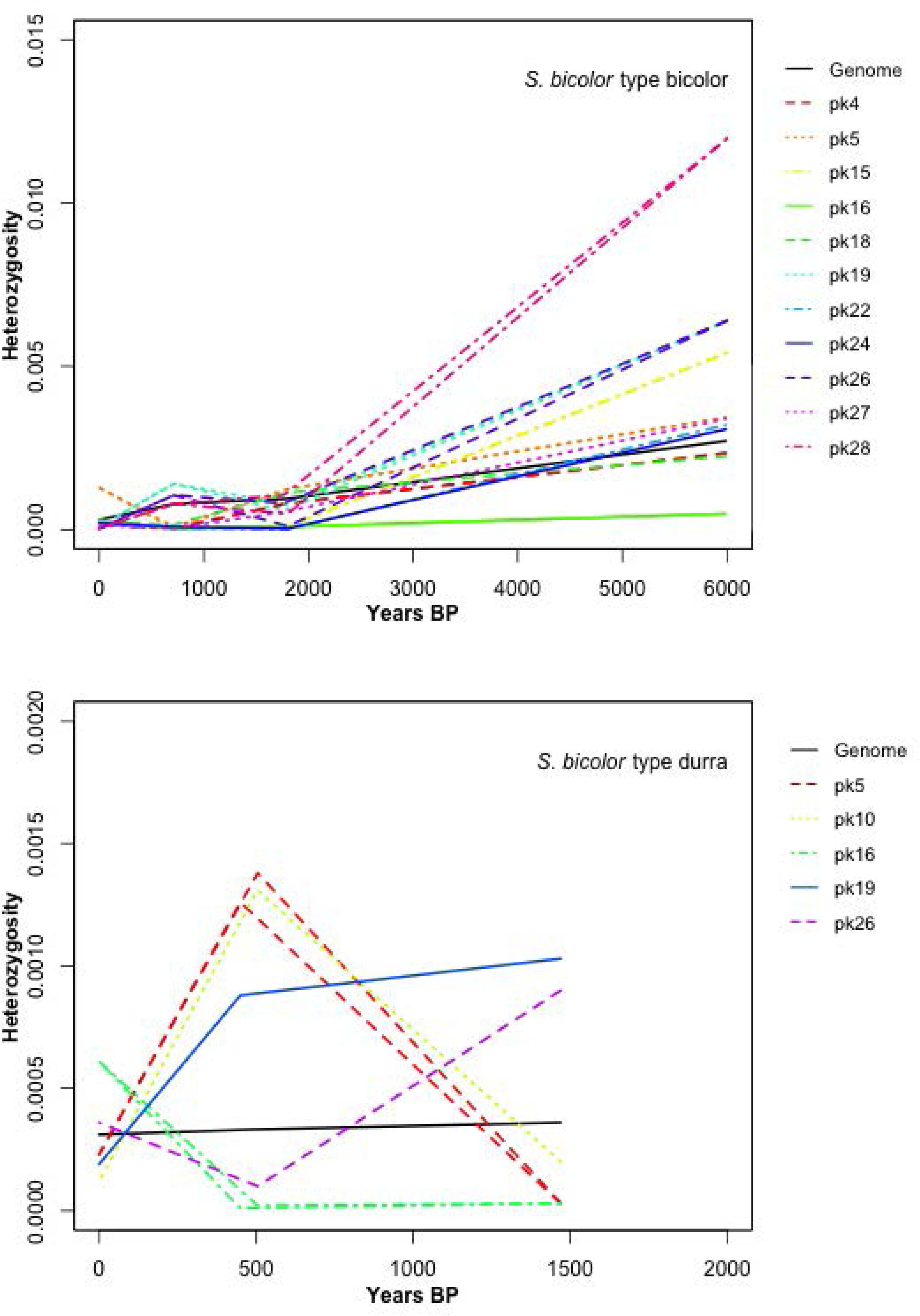
Heterozygosity over time of regions containing high SweeD scores. Significant deviations from the genomic gradient of change over time shown only.

**Figure S7.**
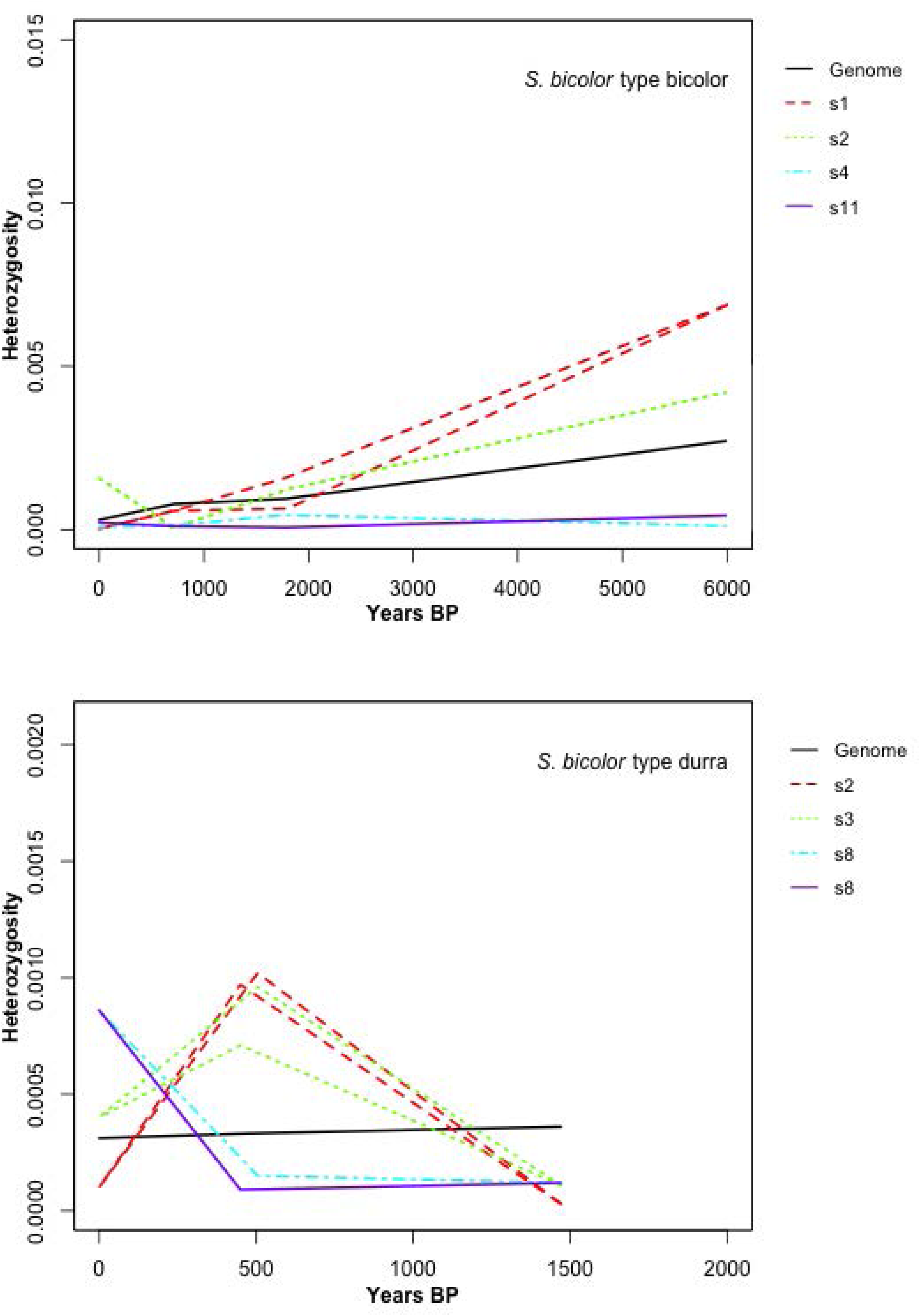
Heterozygosity over time of regions containing domestication loci that have significantly reduced in heterogygosity relative to wild. Significant deviations from the genomic gradient of change over time shown only.

**Figure S8.**
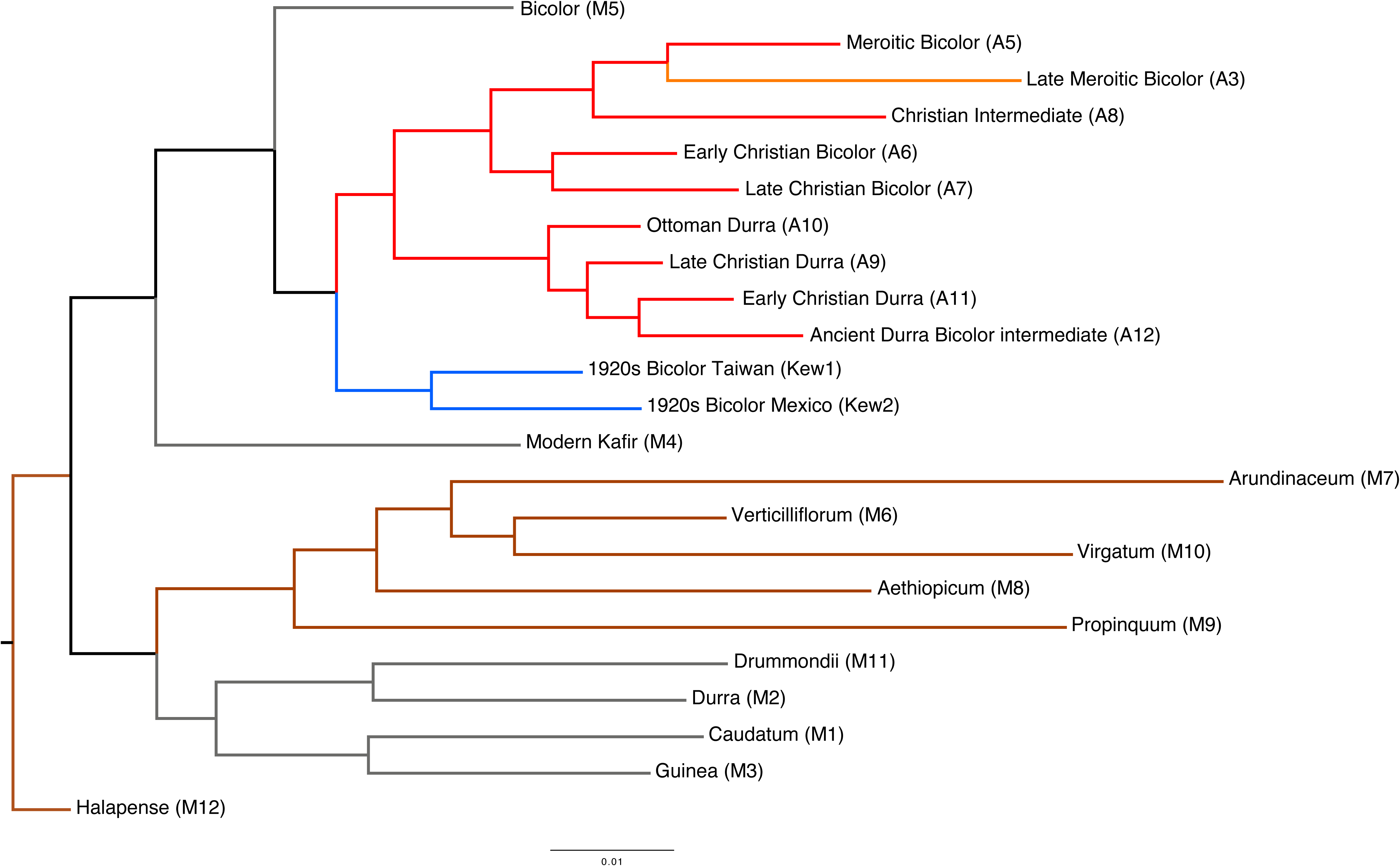
Maximum likelihood tree of whole genome sequence built in EXaML

**Figure S9.**
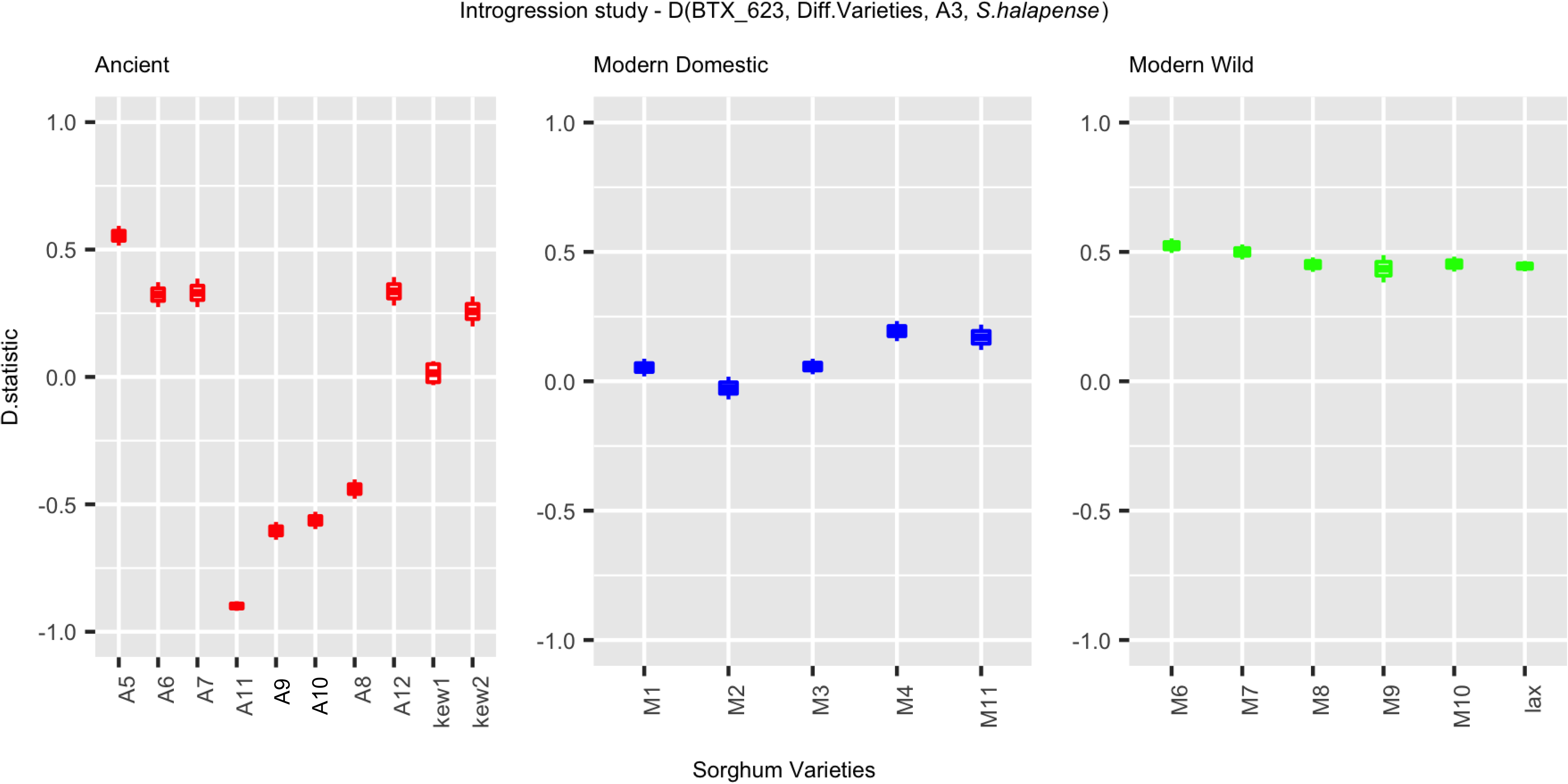
D statistic analysis: P_1_ = *S. bicolor* type bicolor BTX623, P_2_ = sample displayed on X axis, P_3_ = sample A3, P_4_ halapense.

**Figure S10.**
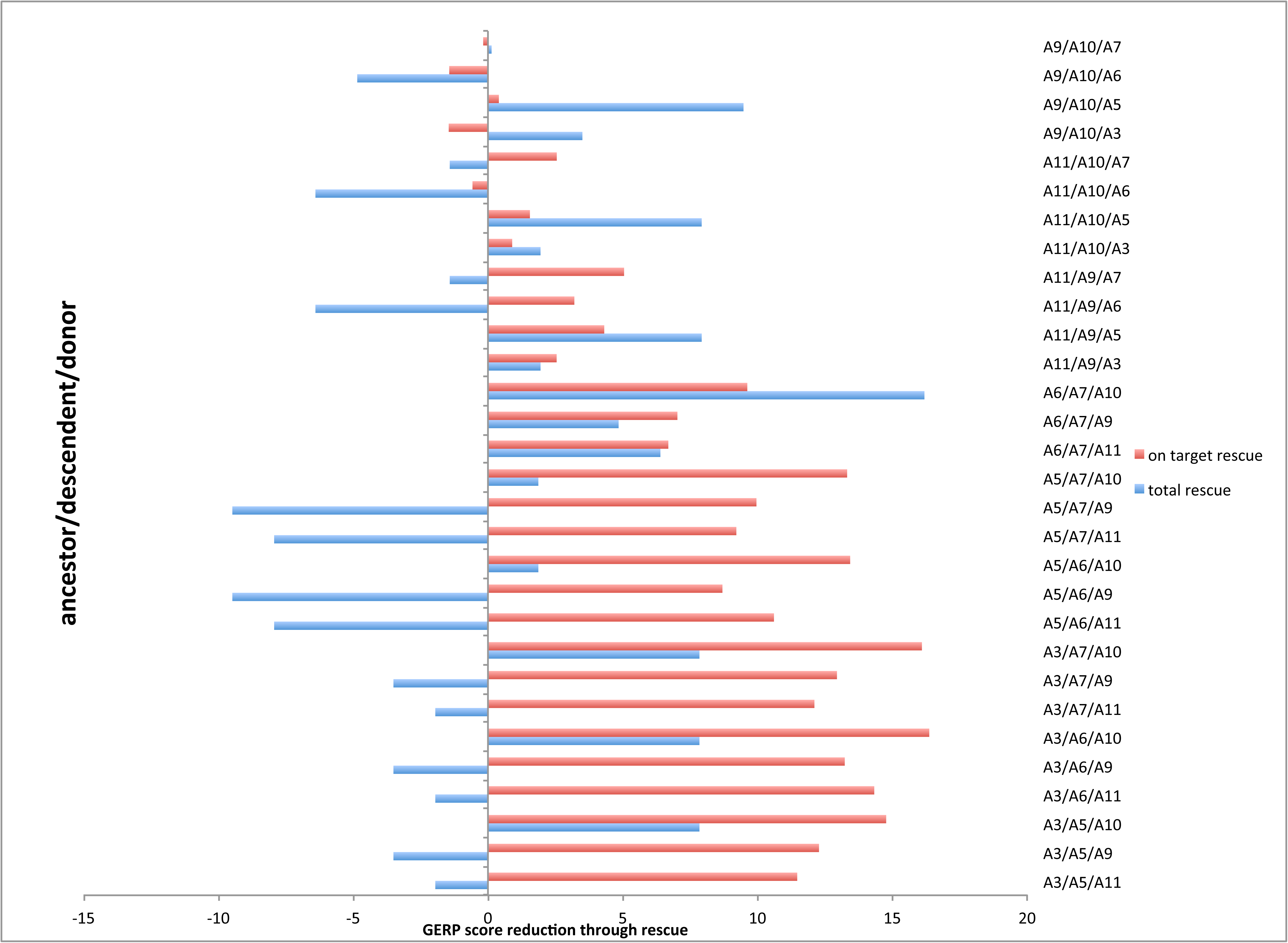
Potential genome rescue of descendents from ancestors by donors based on GERP scores. Red indicates the resultant change from combined score of ancestors and donors in regions of observed GERP load reduction in descendents. Blue indicates the genome wide change in gerp score from combining ancestor and donor scores.

**Figure S11.**
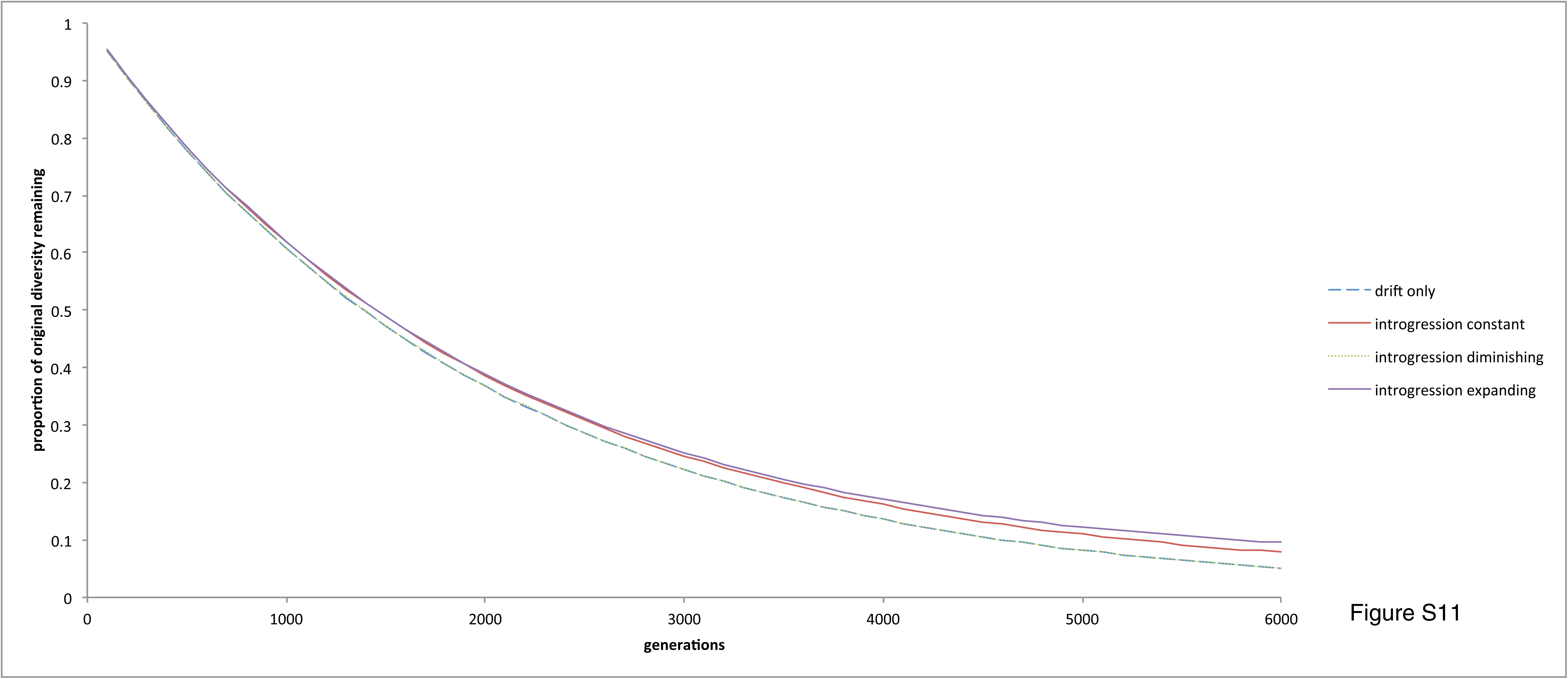
Standard model of loss of genetic diversity through drift combined with introgression over time. Arbitrary founding population of 2000 individuals simulated for 6000 generations to match the over all decrease observed in sorghum. Four models considered, no introgression (drift only), constant introgression (adding 0.000015 to the genetic diversity each generation). Dynamic introgression was defined where the gene flow (gf) contribution each generation is *gf*^*tf*^, where *t* is the generation number and *f* the modification factor. Diminishing introgression, *f* is 1.0001, increasing introgression *f* is 0.99995. See methods for details of calculations.

**Figure S12.**
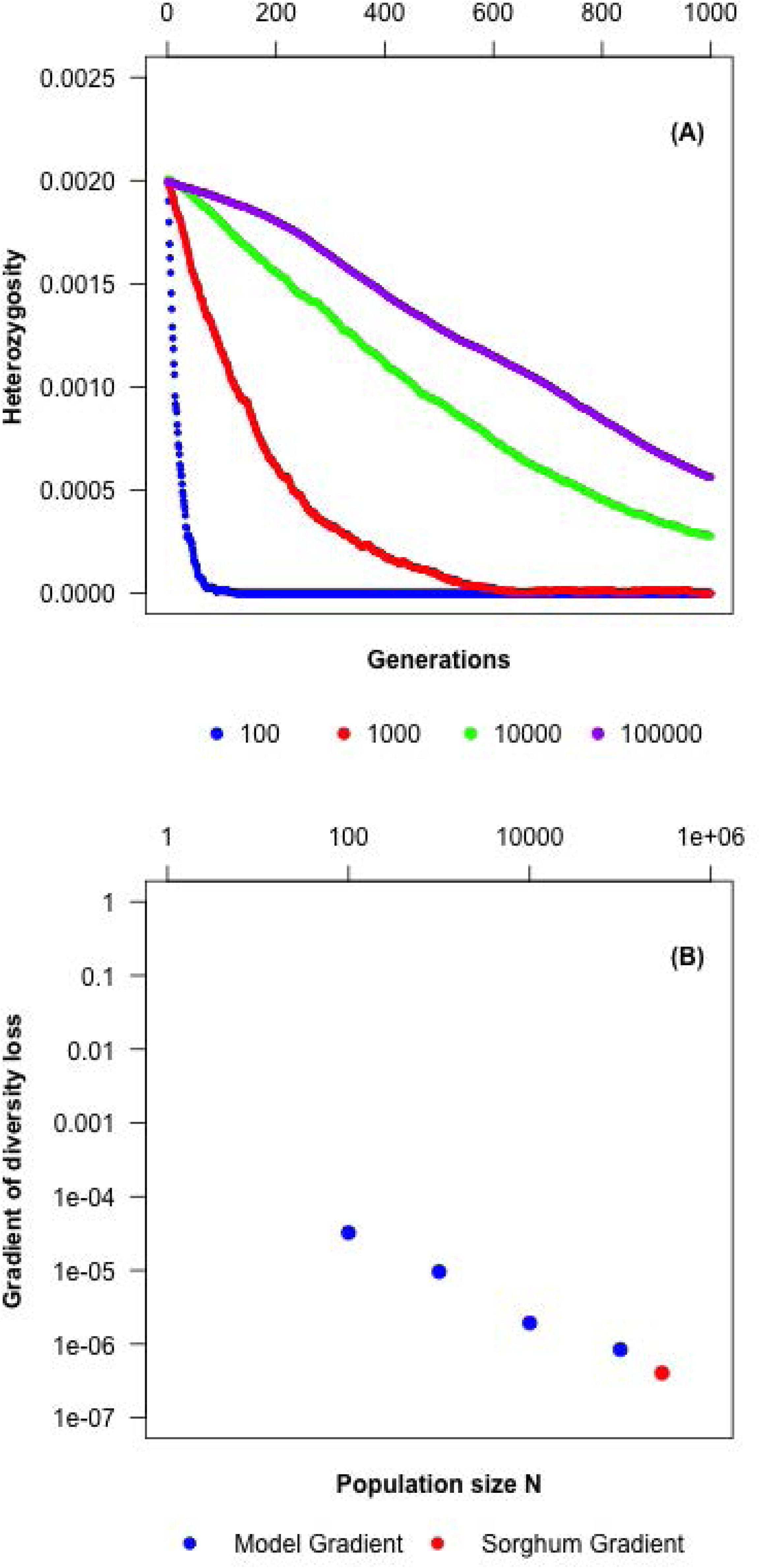
Lost of heterozygosity through founder event model based on crop cycling. A. Sequential founding episodes based on 25% of harvest set aside for sowing (28) for various populations sizes (N). B. Gradients of diversity loss over time in the model and the observed gradient in sorghum with N obtained through linear regression of the model outputs.

**Table S1.**
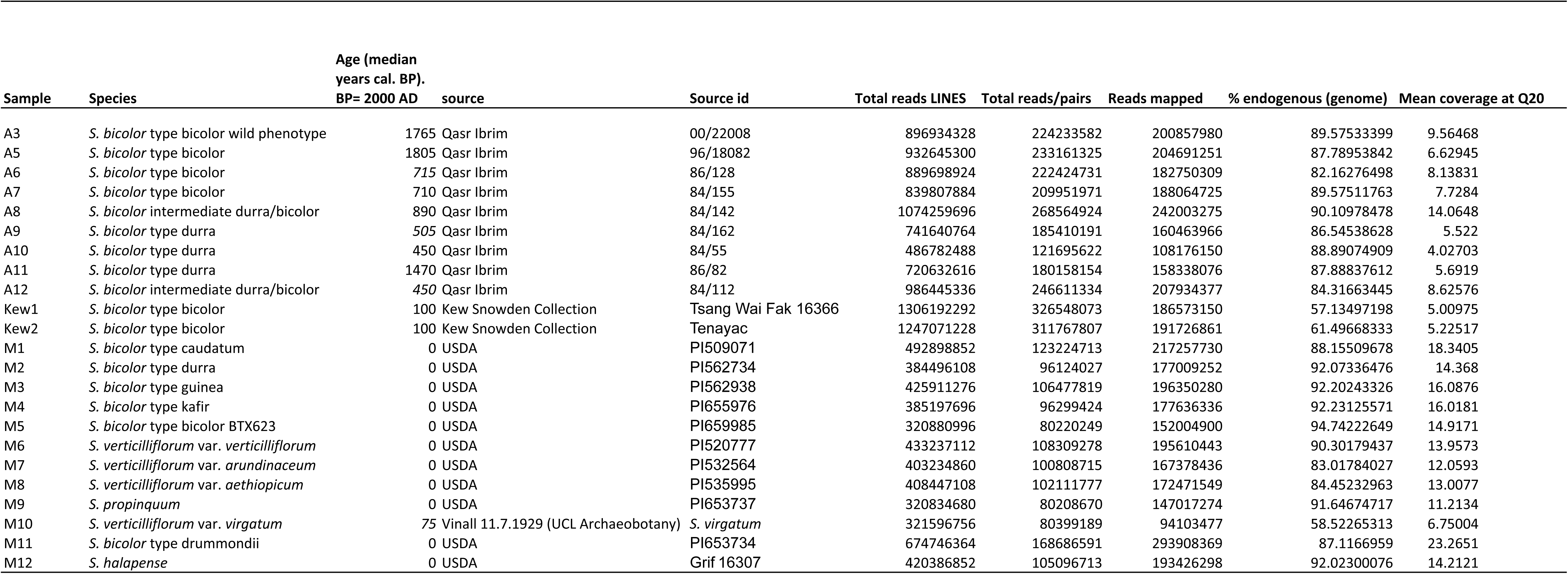
Sample details

**Table S2.**
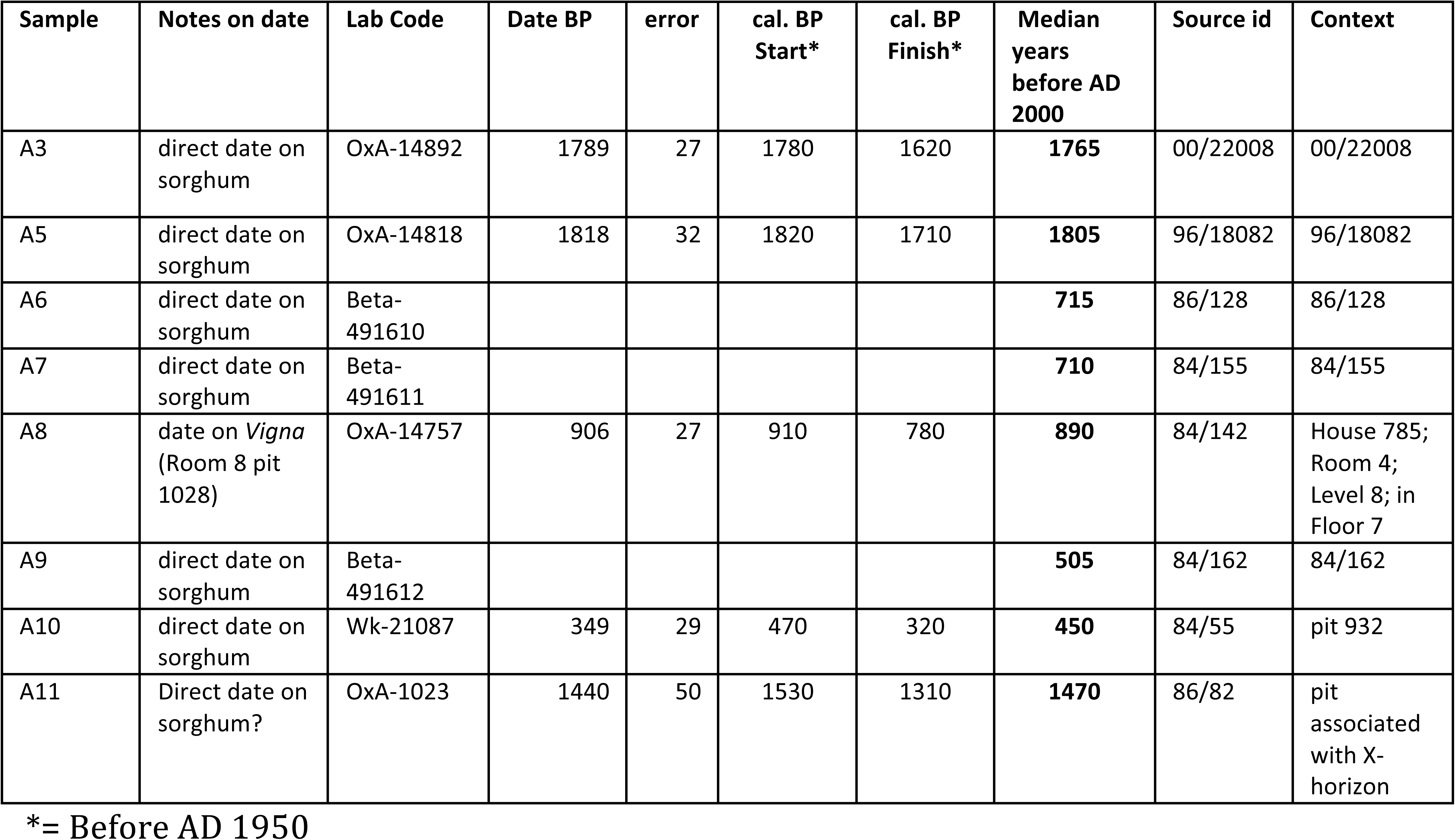
Radiocarbon dates on sorghum specimens or closely associated plant remains

**Table S3.**
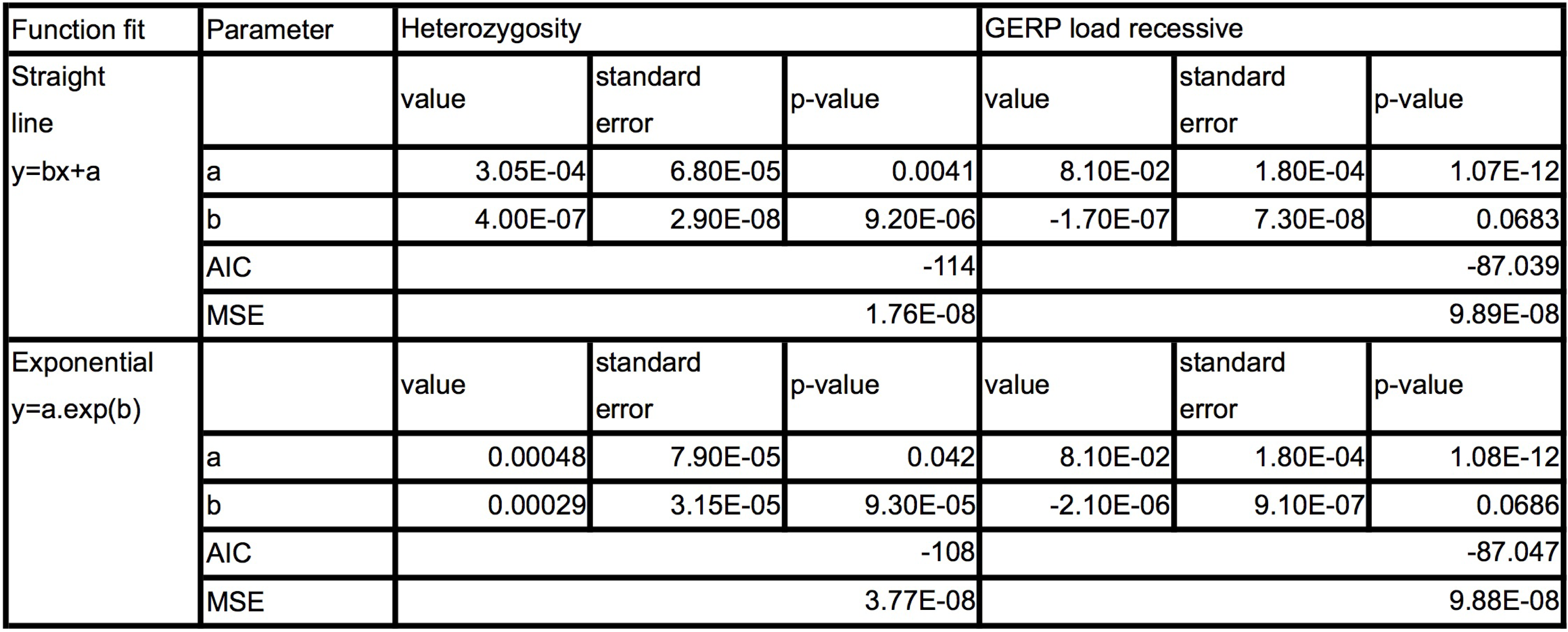
parameter values for curves fit to bicolor heterozyg gosity and GERP data (versus Years BP)

**Table S4.**
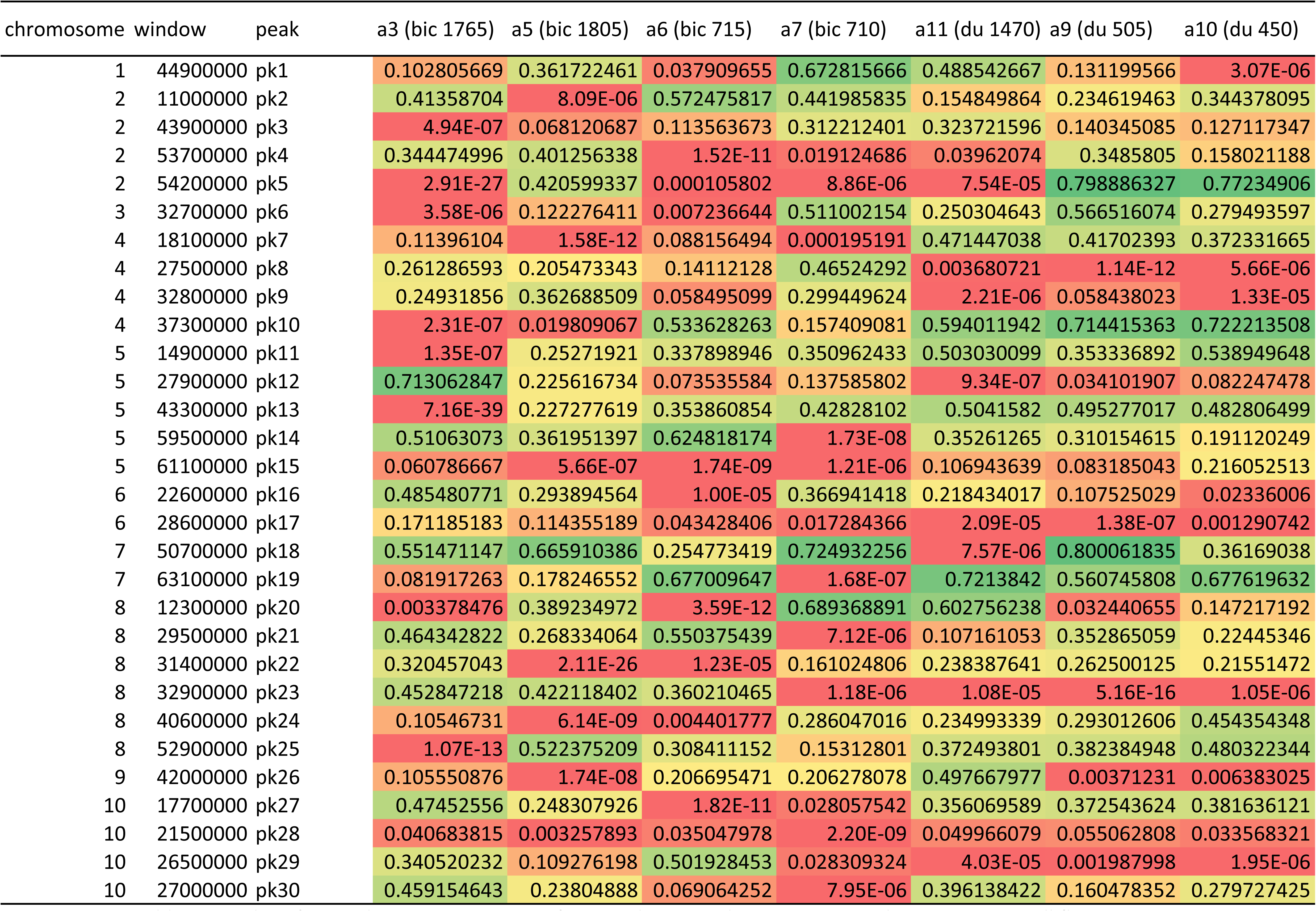
*p* values for windows containing significant reduction in heterozygosity relative to *S. verticilliflorum*

**Table S5.**
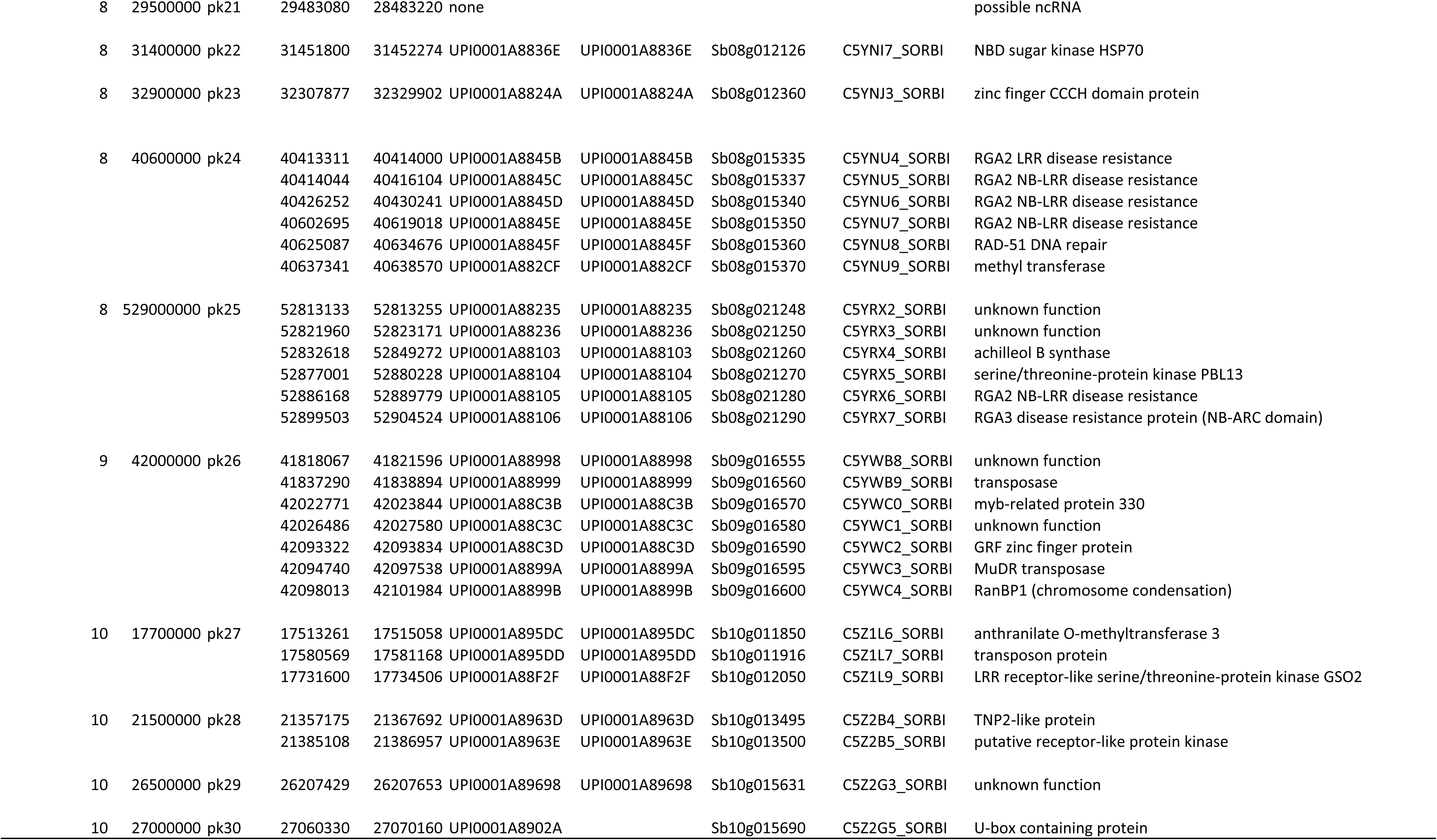
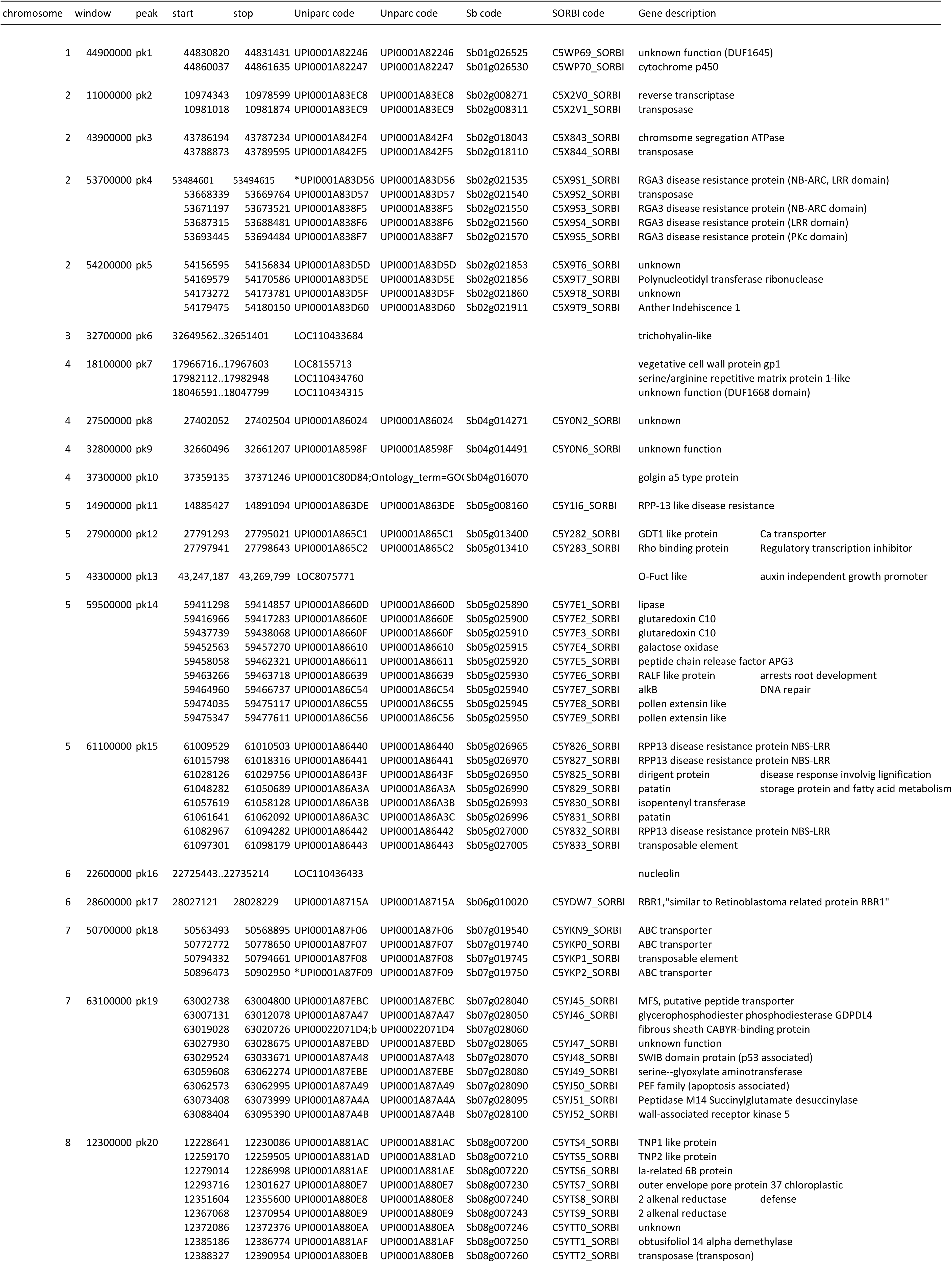
Regions of genome-wide significance in reduction of heterozygosity relative to *S. verticilliflorum* and associated genes

**Table S6.**
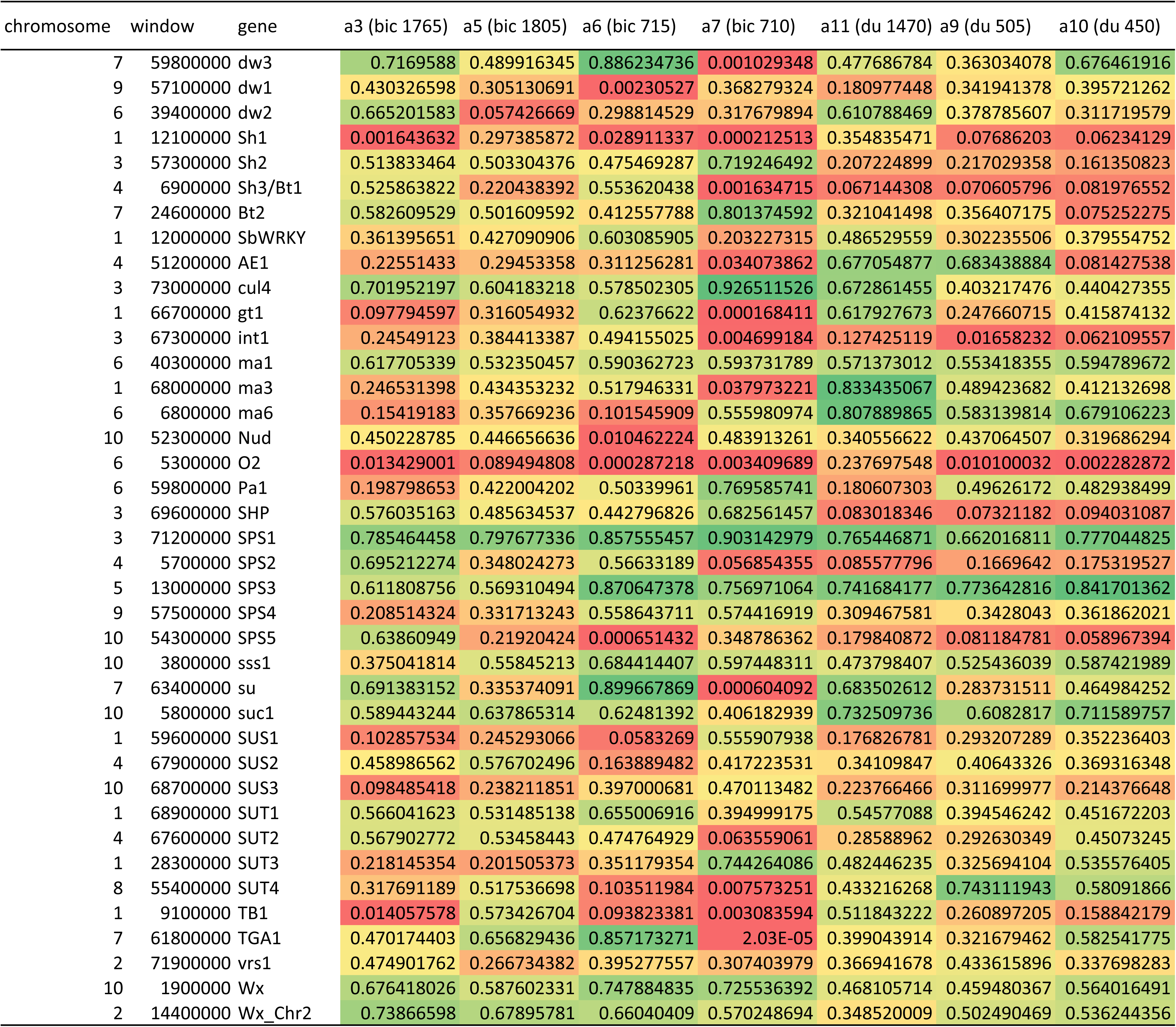
*p* values for reduction in heterozygosity in windows containing domestication loci observed in archaeological accessions relative to *S. verticilliflorum*.

**Table S7.**
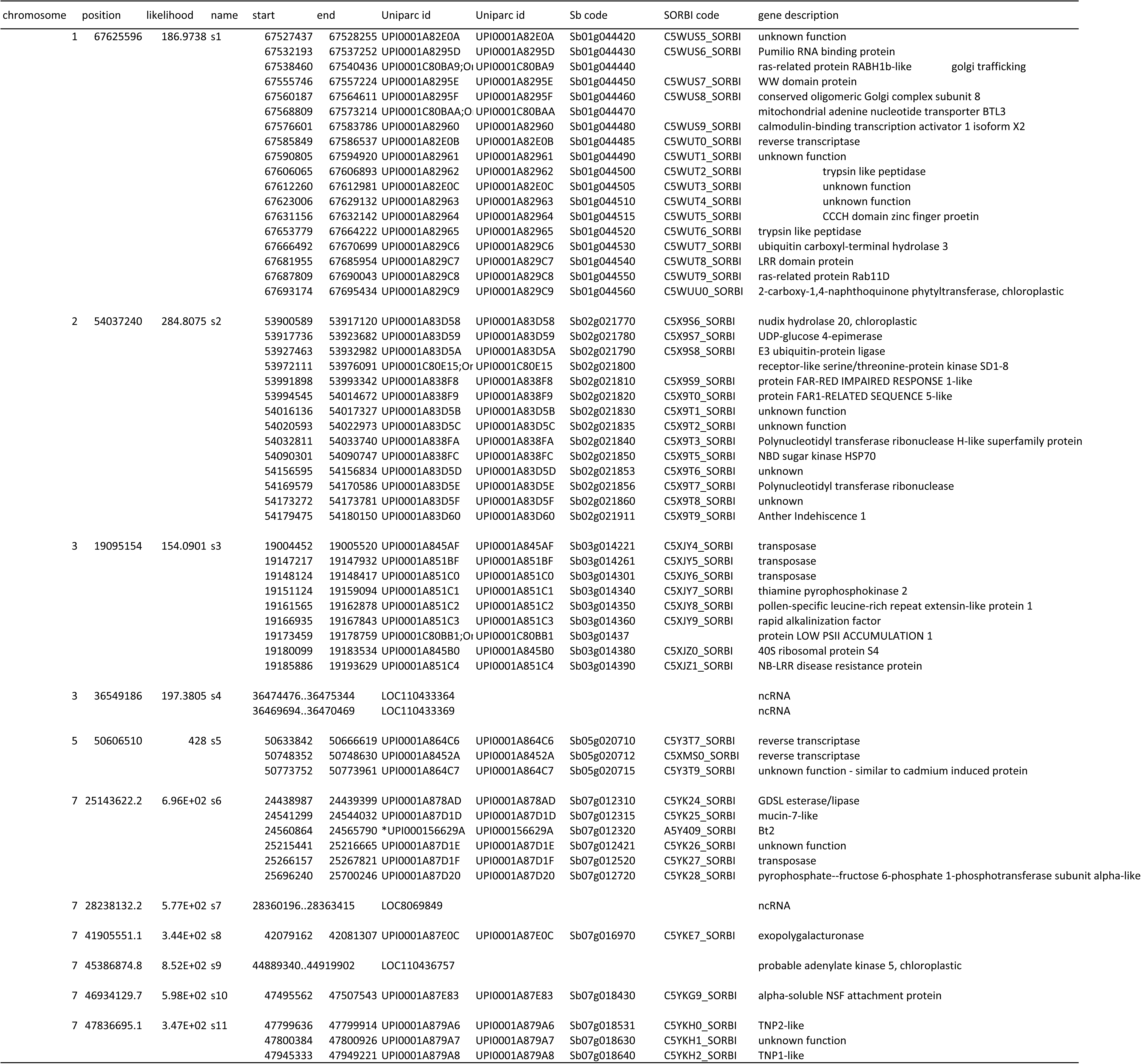
Selective sweep regions identified with SweeD and associated genes * indicates correspondance with the Mace et al (24) candidate domestication gene list

**Table S8.**
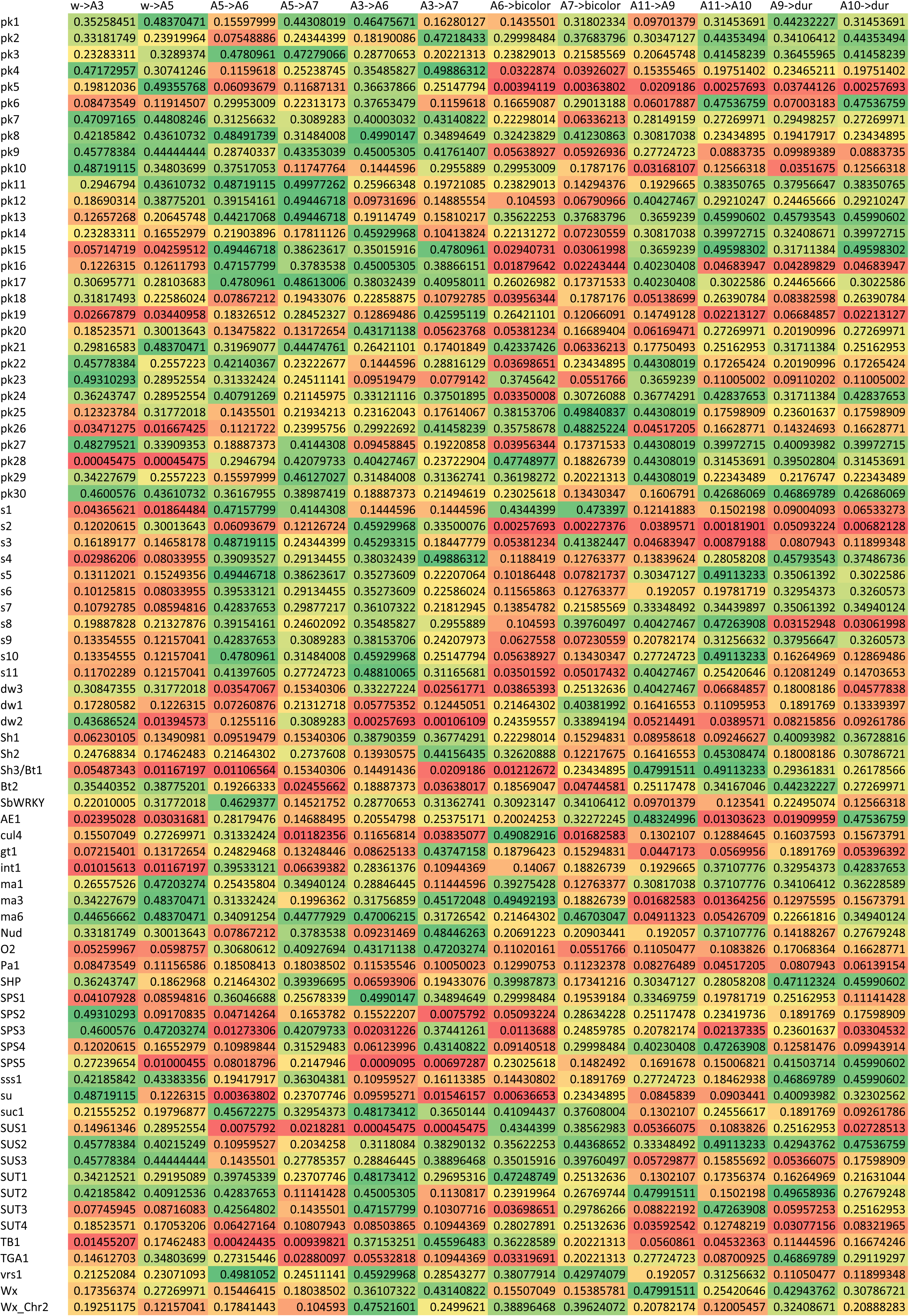
Probabilities of gradients for particular sample transitions against genomic average

**Table S9.**
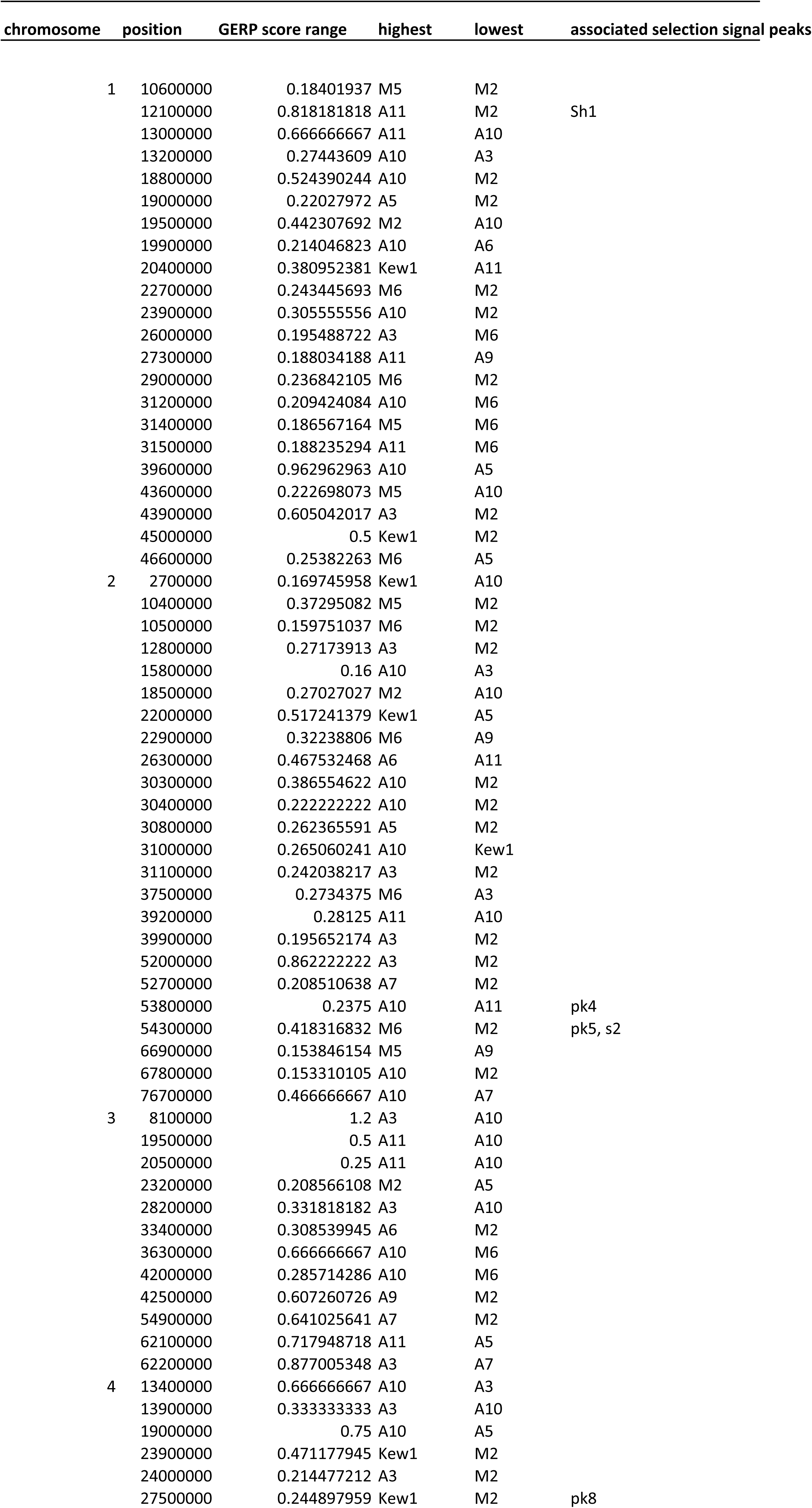
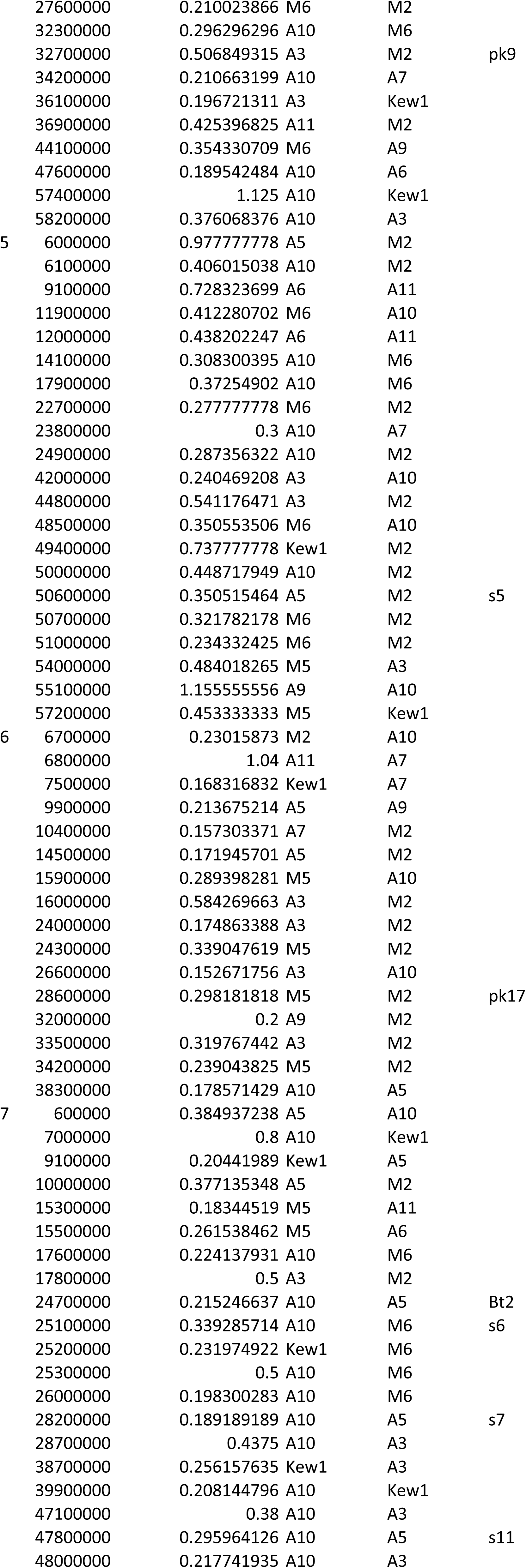
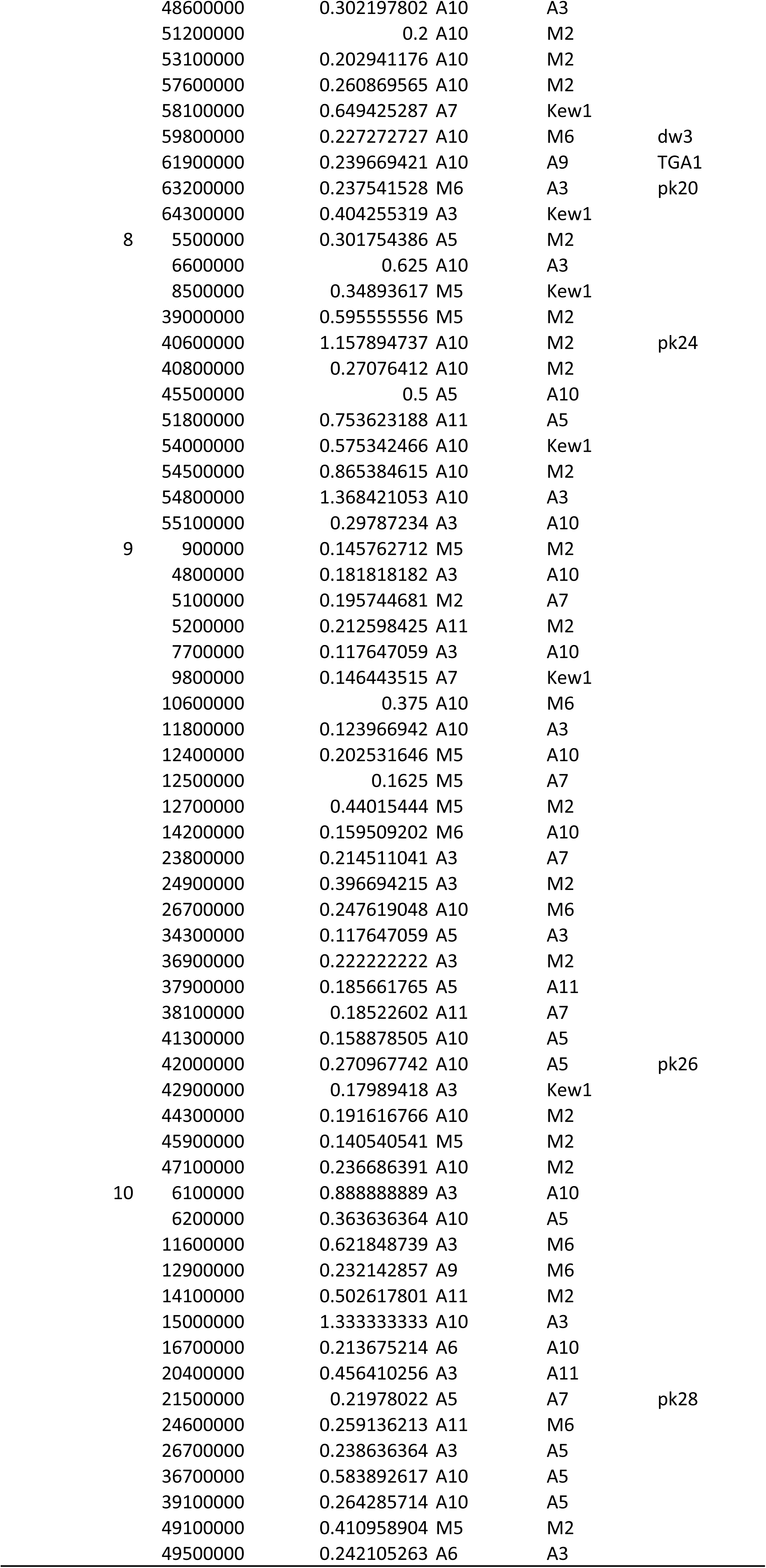
Regions of GERP score deviation between genomes > 2 standard deviations

**Table S10.**
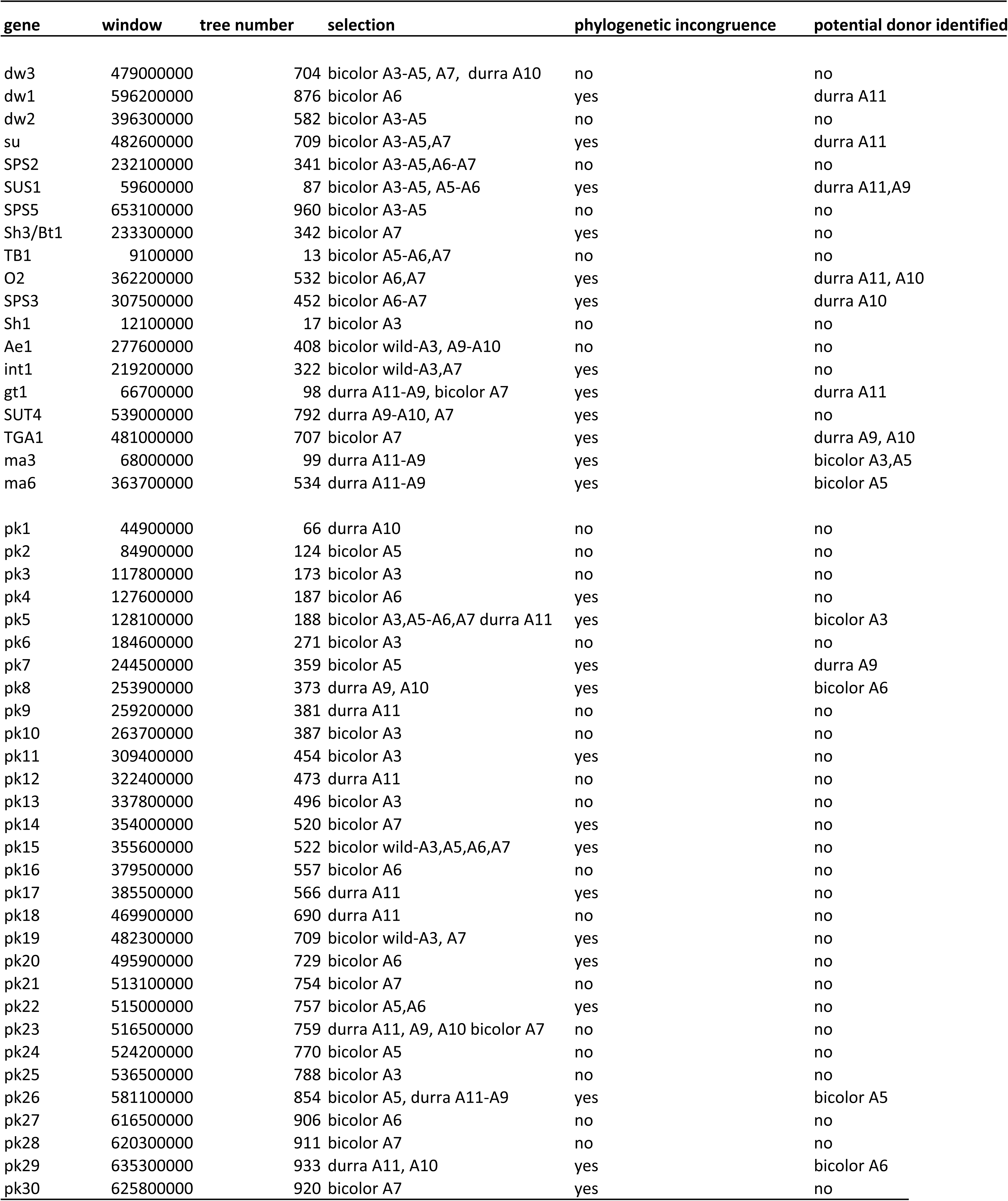
Phylogenetic congruence of regions containing significant reductions in heterozygosity

## Reference

(1) Larson G, Piperno D, Allaby RG et al. Current perspectives and the future of domestication studies. Proc. Natl. Acad. Sci. U.S.A. 111, 6139–6146 (2014).

(2) Purugganan MD, Fuller DQ. The nature of selection during plant domestication. Nature 457, 843–848 (2009).

(3) Fuller DQ, Denham T, Arroyo-Kalin M, Lucas L, Stephens C, Qin L, Allaby RG, Purugganan MD Convergent evolution and parallelism in plant domestication revealed by an expanding archaeological record. Proc. Natl. Acad. Sci. U.S.A. 111, 6147–6152 (2014).

(4) Allaby RG, Stevens S, Lucas L, Maeda O, Fuller DQ. Geographic mosaics and changing rates of cereal domestication. Philosophical Transactions of the Royal Society B. 372, 20160429 (2017).

(5) Poets AM, Fang Z, Clegg MT, Morell PL. Barley landraces are characterized by geographically heterogeneous genomic origins. Genome Biology 16, 173 (2015).

(6) Food and Agriculture Organization of the United Nations. [http://www.fao.org/index_en.htm].

(7) Winchell F, Stevens CJ, Murphy C, Champion L, Fuller DQ. Evidence for Sorghum Domestication in Fourth Millennium BC Eastern Sudan: Spikelet Morphology from Ceramic Impressions of the Butana Group. Current Anthropology 58, 673–683 (2017).

(8) Doggett H Sorghum 2^nd^ Ed, Longman, Harlow (1988).

(9) Brown PJ, Myles S, Kresowich S. Genetic support for a phenotype-based racial classification in sorghum. Crop Sci. 51, 224–230 (2011).

(10) Morris G, et al. Population genomic and genome-wide association studies of agroclimatic traits in sorghum. Proc. Natl. Acad. Sci. U.S.A. 110, 453–458 (2013).

(11) Fuller, Dorian Q and Chris J. Stevens (n.d.) Sorghum Domestication and Diversification: A current archaeobotanical perspective. In: Anna Maria Mercuri, A. Catherine D’Andrea, Rita Fornaciari, Alexa Höhn (eds.) Plants and People in Africa’s Past. Progress in African Archaeobotany. Springer (2017).

(12) Clapham AJ, Rowley-Conwy PA In Fields of Change–Progress in African Archaeobotany, Cappers R, ed. Groningen Archaeological Studies. Groningen, 5, 157–164 (2007).

(13) de Wet JML, Harlan JR, Price EG Variability in *Sorghum bicolor*. In: Harlan JR, de Wet JMJ, Stemler ABL (eds) Origins of African plant domestication. Mouton Press, The Hague, p 453–463 (1976).

(14) Harlan JR, Stemler ABL The races of Sorghum in Africa. In: Harlan JR, de Wet J, Stemler ABL (eds) Origins of African plant domestication. Mouton Press, The Hague, p 465–478 (1976).

(15) Meyer R, Purugganan M. Evolution of crop species: genetics of domestication and diversification. Nature Reviews Genetics 14, 840–852 (2013).

(16) Liu Q, Zhou Y, Morrell PL, Gaut BS Deleterious variants in Asian rice and the potential cost of domestication. Mol. Biol. Evol. 34(4), 908–924 (2017).

(17) Renault S, Rieseberg L. The accumulation of deleterious mutations as a consequence of domestication and improvement in sunflower and other Compositae crops. Mol. Biol. Evol. 32(9), 2273–2283 (2015).

(18) Wang L, Beissinger TM, Lorant A, Ross-Ibarra C, Ross-Ibarra J, Hufford MB The interplay of demography and selection during maize domestication and expansion. Genome Biol. 18, 215 (2017).

(19) Cooper GM et al. Distribution and intensity of constraint in mammalian genomic sequence. Genome Res. 15, 901–913 (2005).

(20) Smith O, Clapham A, Rose P, Liu Y, Wang J, Allaby RG. Genomic methylation patterns in archaeological barley show de-methylation as a time-dependent diagenetic process. Scientific Reports 4, 5559 (2014).

(21) Pavlidis P, Živkovic D, Stamatakis A, Alachiotis N SweeD: Likelihood-based detection of selective sweeps in thousands of genomes. Mol Biol Evol 30, 2224–2234 (2013).

(22) Hudson M, Ringli C, Boylan MT, Quail PH The *FAR1* locus encodes a novel nuclear protein specific to phytochrome A signaling. Genes Devel. 13, 2017–2027 (1999).

(23) Zhu QH, Ramm K, Shivakkumar R, Dennis ES, Upadhyahya NM The *ANTHER INDEHISCENCE1* gene encoding a single MYB domain protein is involved in anther development in rice. Plant Physiol. 135, 1514–1525 (2004).

(24) Martin, S, Davey, J, Jiggins, C. Evaluating the Use of ABBA–BABA Statistics to Locate Introgressed Loci. Mol. Biol. Evol. 32(1), 244–257 (2014).

(25) Mace ES et al. Whole genome sequencing reveals untapped genetic potential in Africa’s indigenous cereal crop sorghum. Nature Communications 4, 2320 (2013).

(26) Thurber CS, Ma JM, Higgins RH, Brown PJ Retrospective genomic analysis of sorghum adaptation to temperate-zone grain production. Genome Biol. 14, R68 (2013).

(27) DeGiorgio M, Jakobsson M, Rosenberg N. Explaining worldwide patterns of human genetic variation using a coalescent-based serial founder model of migration outward from Africa. Proc. Natl. Acad. Sci. U.S.A. 106: 16057–16062 (2009).

(28) Hillman GC, Davies MS. Domestication rates in wild type wheats and barley under primitive cultivation. B. J. Linn. Soc. 39:39–78 (1990).

(29) Kremling KA et al. Dysregulation of expression correlates with rare-allele burden and fitness loss in maize. Nature 555:520–523 (2018).

(30) Rogers & Slatkin Excess defects in a woolly mammoth on Wrangel Island PloS Genetics 13(3): e1006601 (2017).

(31) Shennan S, Downey SS, Timpson A, et al. Regional population collapse followed initial agricultural booms in mid–Holocene Europe. Nat Commun. 4:2486 (2013).

(32) Allaby RG, Kitchen JL, Fuller DQ (2016) Surprisingly low limits of selection in plant domestication. Evolutionary Bioinformatics 11:(S2) 41–51.

(33) Alexander, J The Saharan divide in the Nile Valley: the evidence from Qasr Ibrim. The African Archaeological Review 6, 73–90 (1988).

(34) Rose P (2013) Qasr Ibrim In: Bagnall RS, Brodersen K, Champion CB, Erskine A, Huebner SR (eds.) The Encyclopedia of Ancient History. Oxford: Wiley. Pp. 5695–5697 DOI: 10.1002/9781444338386.wbeah15340.

(35) Palmer SA, Moore JD, Clapham AJ, Rose P, Allaby RG. Archaeogenetic evidence of ancient Nubian barley evolution from six to two-row indicates local adaptation. PLoS One, 4(7), e6301 (2009).

(36) Palmer SA, Clapham AJ, Rose P, Freitas F, Owen BD, Beresford-Jones D, Moore JD, Kitchen JL, Allaby RG. Archaeogenomic evidence of punctuated genome evolution in Gossypium. Molecular biology and evolution, 29(8), 2031–2038 (2012).

(37) O’Donoghue K, Clapham A, Evershed RP, Brown TA. Remarkable preservation of biomolecules in ancient radish seeds. Proc R Soc B Biol Sci. 263, 541–547 1996.

(38) Alexander, J, Driskell B. Qasr Ibrim 1984. Journal of Egyptian Archaeology 71, 12–26 (1985).

(39) Rowley-Conwy P Nubia AD 0-550 and the ‘Islamic’ agricultural revolution: Preliminary botanical evidence from Qasr Ibrim, Egyptian Nubia. Archeologie du Nil Moyen 3, 131–138 (1989).

(40) Clapham AJ, Rowley-Conwy PA (2007). New discoveries at Qasr Ibrim, Lower Nubia. Fields of change: progress in African archaeobotany. Barkhuis & Groningen University Library, Groningen, The Netherlands, 157–164.

(41) Clapham, A., & Rowley-Conwy, P. (2009). The archaeobotany of cotton (Gossypium sp. L.) in Egypt and Nubia with special reference to Qasr Ibrim, Egyptian Nubia. From foragers to farmers. Papers in honour of Gordon C. Hillman. Oxbow Books, Oxford, 244–253.

(42) Adams, WY (1986) Ceramic Industries of Medieval Nubia. University of Kentucky Press

(43) Hanghøj K, Seguin-Orlando A, Schubert M, Madsen T, Pedersen JS, Willerslev E, Orlando L. Fast, Accurate and Automatic Ancient Nucleosome and Methylation Maps with epiPALEOMIX. Mol. Biol. Evol. 33, 3248–3298 (2016).

(44) Paterson AH et al. The Sorghum bicolor genome and the diversification of grasses. Nature 457, 551–556 (2009).

(45) Bouyer et al. DNA methylation dynamics during early plant life. Genome Biology 18:179 (2017)

(46) Langmead B, Salzberg SL. Fast gapped-read alignment with Bowtie 2. Nature Methods. 9(4), 357–359 (2012).

(47) Li H, Handsaker B, Wysoker A, Fennell T, Ruan J, Homer N, Marth G., Abecasis G., Durbin R. and 1000 Genome Project Data Processing Subgroup The Sequence alignment/map (SAM) format and SAMtools. Bioinformatics 25, 2078–2079 (2009).

(48) da Fonseca RR, Smith BD, Wales N, Cappellini E, Skoglund P, Fumagalli M, Samaniego JA, Carøe C, Ávila-Arcos MC, Hufnagel DE, Korneliussen TS, Vieira FG, Jakobsson M, Arriaza B, Willerslev E, Nielsen R, Hufford MB, Albrechtsen A, Ross-Ibarra J, Gilbert MT The origin and evolution of maize in the Southwestern United States. Nature Plants 1, 14003 (2015).

(49) Koslov AM, Aberer A, Alexandros S. ExaML version 3: a tool for phylogenomic analysis on supercomputers. Bioinformatics 31, 2577–2579 (2015).

(50) Felsenstein J Evolutionary trees from DNA sequences: A maximum likelihood approach. J Mol Evol 17(6), 368–376 (1981).

(51) Stamatakis A RAxML-VI-HPC: maximum likelihood-based phylogenetic analyses with thousands of taxa and mixed models. Bioinformatics 22(21), 2688–2690 (2006).

(52) Van der Auwera GA, Carneiro M, Hartl C, Poplin R, del Angel G, Levy-Moonshine A, Jordan T, Shakir K, Roazen D, Thibault J, Banks E, Garimella K, Altshuler D, Gabriel S, DePristo M. From FastQ Data to High-Confidence Variant Calls: The Genome Analysis Toolkit Best Practices Pipeline. Current Protocols In Bioinformatics 43, 11.10.1-11.10.33 (2013).

(53) Pfeifer B, Wittelsbürger U, Ramos-Onsins S, Lercher M. PopGenome: An Efficient Swiss Army Knife for Population Genomic Analyses in R. Mol. Biol. Evol. 31(7):1929–1936 (2014).

(54) Wolfe KH, Sharpe PM, Li WH. Rates of synonymous substitution in plant nuclear genes. J. Mol. Evol. 29:208–211 (1989).

